# Rethinking phylogenetic comparative methods

**DOI:** 10.1101/222729

**Authors:** Josef C. Uyeda, Rosana Zenil-Ferguson, Matthew W. Pennell

## Abstract

As a result of the process of descent with modification, closely related species tend to be similar to one another in a myriad different ways. In statistical terms, this means that traits measured on one species will not be independent of traits measured on others. Since their introduction in the 1980s, phylogenetic comparative methods (PCMs) have been framed as a solution to this problem. In this paper, we argue that this way of thinking about PCMs is deeply misleading. Not only has this sowed widespread confusion in the literature about what PCMs are doing but has led us to develop methods that are susceptible to the very thing we sought to build defenses against — unreplicated evolutionary events. Through three Case Studies, we demonstrate that the susceptibility to singular events is indeed a recurring problem in comparative biology that links several seemingly unrelated controversies. In each Case Study we propose a potential solution to the problem. While the details of our proposed solutions differ, they share a common theme: unifying hypothesis testing with data-driven approaches (which we term “phylogenetic natural history”) to disentangle the impact of singular evolutionary events from that of the factors we are investigating. More broadly, we argue that our field has, at times, been sloppy when weighing evidence in support of causal hypotheses. We suggest that one way to refine our inferences is to re-imagine phylogenies as probabilistic graphical models; adopting this way of thinking will help clarify precisely what we are testing and what evidence supports our claims.

## Introduction

Every so often, evolution comes up with something totally new and unexpected, a so-crazy-it-just-might-work set of adaptations that is the stuff of nature documentaries. Many biologists likely have a favorite example of a lineage that has evolved something spectacular such as devilishly horned lizards that squirt blood from their eye sockets or marine sloths that grazed ancient seabeds.

As macroevolutionary researchers, it is hard to know what to do with these types of events (Vermeij, 2006). Their singular and unreplicated nature seems incompatible with models that we typically use to describe change over time, such as Brownian motion (BM; Felsenstein, 1973) or the Mk model (Pagel, 1994; Lewis, 2001). Such models presume continuity, whereas one-off events, such as the evolution of novel nutritive function in exocrine glands leading to mammalian milk, have no clear precedent in history. The evolution of such traits may set in motion a cascade of changes across an organism, such that descendant lineages may look very different in many ways from their more distant relatives. Or alternatively, a suite of traits may just happen to change at the same time. In either case, it is these sorts of idiosyncratic and unreplicated events that we often think of when we think of the need to consider phylogeny in analyses of comparative data. And this is not an abstract concern; a wide breadth of macroevolutionary data suggest that abrupt shifts and discontinuities have been a major feature of life on Earth (Uyeda et al., 2011, 2017; Landis and Schraiber, 2017; Jablonski, 2017). But as recent controversies in phylogenetic comparative biology have highlighted, our current methods (reviewed in Pennell and Harmon, 2013; O’Meara, 2012; Garamszegi, 2014) are not designed to deal with such dynamics.

For example, Maddison and FitzJohn (2015) recently demonstrated that common statistical tests (e.g., Pagel, 1994; Maddison, 1990) for the evolutionary correlation of discrete characters are prone to reporting a significant association even when the pattern is driven by a single (or, very few) independent transition(s) from one character state to another. Maddison and FitzJohn (2015) referred to such scenarios as cases of ‘phylogenetic pseudoreplication’ (see also Read and Nee, 1995; Nee et al., 1996).

We will argue that this unresolved challenge permeates not just tests for discrete character correlations, but nearly every method of finding associations in comparative methods (Figure 1). For example, Rabosky and Goldberg (2015) show that applying trait-dependent diversification models (e.g., BiSSE; Maddison et al., 2007) to real-world phylogenies, which are usually not shaped like trees resulting from simulations of a birth-death stochastic process (Mooers and Heard, 1997), often leads to support for trait-dependent diversification models regardless of whether traits are actually affecting speciation and extinction. The work of Beaulieu and O’Meara (Beaulieu et al., 2013; Beaulieu and O’Meara, 2014, 2016) has illuminated important underlying reasons behind Rabosky and Goldberg’s findings: the failure to consider alternative models in which the “background” diversification rate changes across the tree (i.e., there is a shift in diversification regimes unrelated to the trait being considered). To address this shortcoming, Beaulieu et al. (2013) borrowed an idea from molecular phylogenetics (Penny et al., 2001; Galtier, 2001), and developed a Hidden States Model (HSM) for describing the evolution of a binary character along a phylogeny. In their HSM the transition rates between character states depend on the ‘hidden’ state of another, unobserved, trait also evolving along the tree (also see Price, 1997; Felsenstein, 2011, both of whom explored a related model). Applying the same principle to trait-dependent diversification models, they showed how models that include background heterogeneity in diversification rates provide a fairer comparison to the hypothesis of genuine state-dependent diversification (Beaulieu and O’Meara, 2016). Rather than considering a biologically unrealistic constant-rate null hypothesis, Beaulieu and colleagues built models that allowed traits and diversification to vary in biologically plausible ways (also see Zenil-Ferguson and Pennell, 2017, on this point).

**Figure 1:**
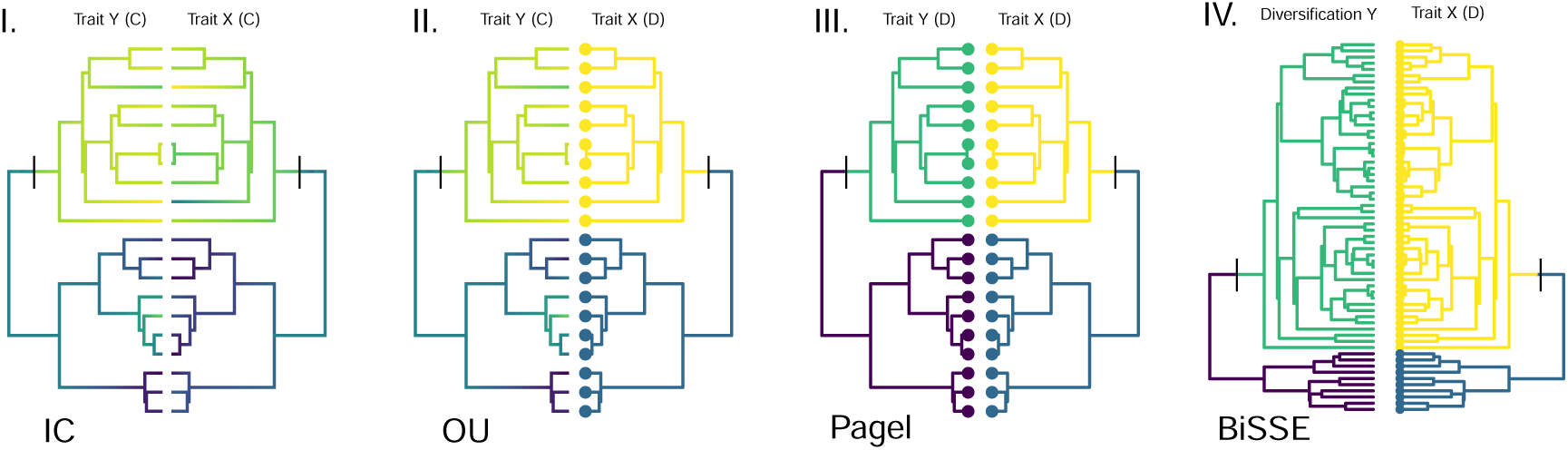
Singular, unreplicated events (vertical dashes) can generate apparently significant associations across several types of comparative analyses. Case Studies I–III are indicated in panels I–III, and though we do not consider diversification models such as BiSSE in our examples, they are similarly affected (panel IV). In each case, we map (in some cases, arbitrarily) the dependent variable (*Y*) on the phylogeny on the left and the predictor trait on the same phylogeny to the right (*X*), and indicate whether the trait is a continuous trait (C), a discrete trait (D) or a diversification rate. We also suggest a common method used to analyze such associations: IC - Independent Contrasts (Felsenstein, 1985); OU - Ornstein-Uhlenbeck models (Butler and King, 2004); Pagel - Pagel’s correlation test (Pagel, 1994). Colors on the branches indicate the state of the character on the phylogeny — either continuous trait value, discrete character state, or diversification rate regime. Panels I and III correspond to variations of “Felsenstein’s worst-case scenario” and “Darwin’s scenario”, respectively.

We think that the solution proposed by Beaulieu and O’Meara (2016) — accounting for background shifts in evolutionary regimes unrelated to the focal trait association— is general and applies across comparative biology. In this paper we develop this argument through a series of three Case Studies, depicted in panels I–III of Figure 1. We will show in each Case Study that rare evolutionary events may deceive our methods and distort our interpretations. For each study, we will then sketch out possible solutions for making causal inferences from comparative data. These solutions differ in their modeling details and methods of inference but they share a core idea.

More specifically, all three Case Studies revolve around the problem of how to discover plausible histories of singular events, or transitions in evolutionary regimes — a practice we call “phylogenetic natural history” — and how to disentangle the impact of these events from that of the hypothesized effects we are investigating. In our examples, we highlight scenarios where a single change in the background evolutionary dynamics can lead to apparent associations between factors of interest, as we find such cases particularly illuminating. But the problems (and potential solutions) we identify apply just as well to situations where the background evolutionary dynamics change more frequently.

By working through the Case Studies, we arrive at two general recommendations for how to move phylogenetic comparative methods (PCMs) forward. First, we advocate for unifying hypothesis-testing and data-driven approaches. Rather than being alternative methods of investigating macroevolutionary processes and patterns, they are complementary, and in our view, essential, to one another. Second, we propose that comparative biologists need to be more careful about how we draw causal inference from phylogenetic data. One particularly elegant solution is to render comparative analyses as graphical models. These graphical models can help clarify exactly what causal statements we are making and what the limits of these inferences are.

### Case Study I: Felsenstein’s worst-case scenario

More than anything else, it was the famous series of figures depicting the “worst-case scenario” (Figures 5, 6, and 7 in the original; our Figure 2) from Felsenstein’s iconic 1985 paper “Phylogenies and the comparative method” that awakened biologists to the need for tree-thinking and started a revolution in modern comparative biology. The idea is simple: as a result of shared ancestry, measurements taken on one species will not be independent from those collected on another and especially so, if the two species are closely related. This non-independence can create apparent correlations between traits that, are in truth, evolving independently. To illustrate the effect of non-independence of characters, Felsenstein generated a scenario in which two clades are separated by long branches (our Figure 2). He then evolved traits according to a BM process along the phylogeny; he recovered a significant regression slope using Ordinary Least Squares (OLS) despite there being no evolutionary covariance between the traits.

**Figure 2:**
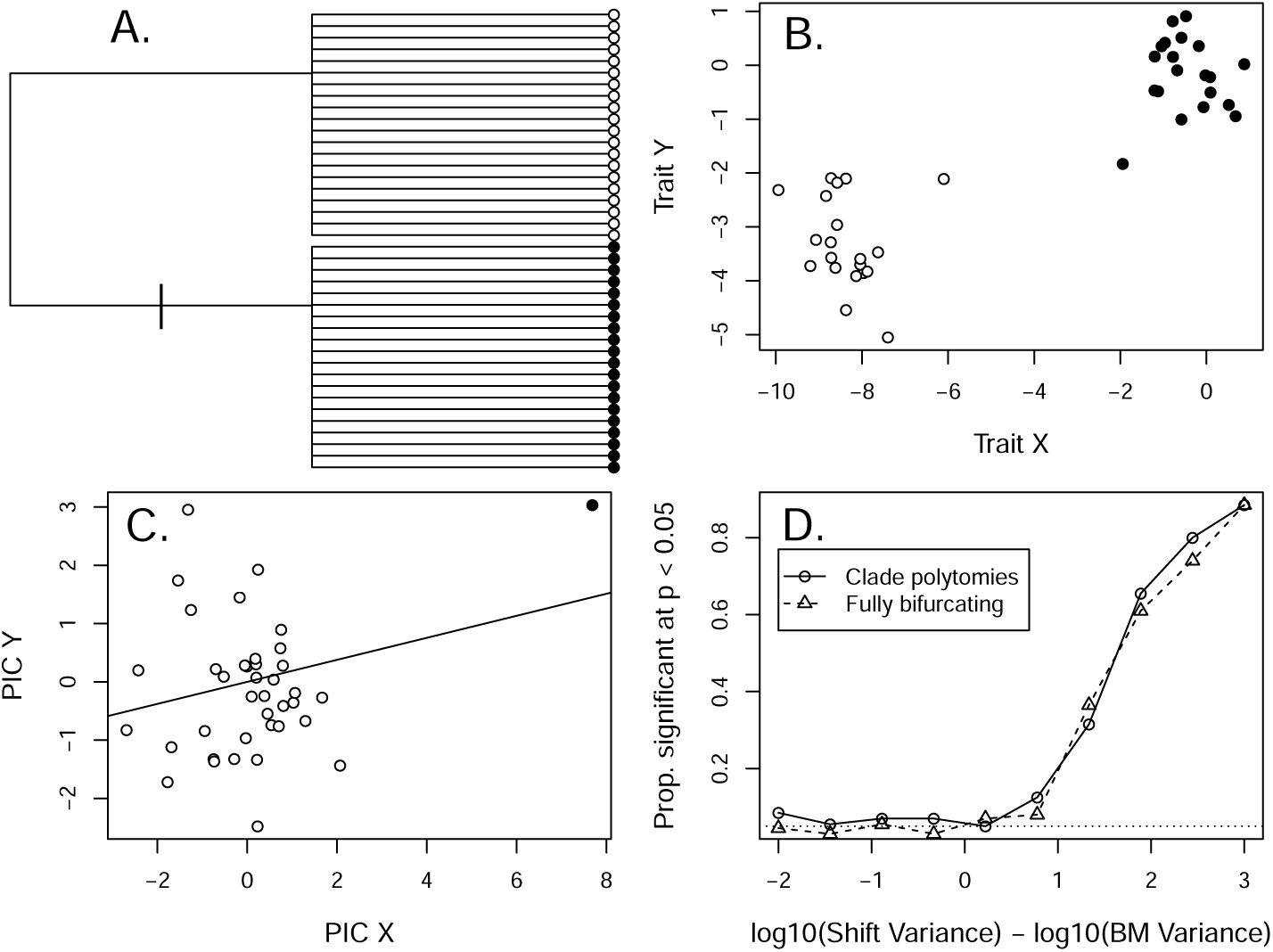
Felsenstein’s worst-case scenario (Felsenstein, 1985) illustrates a problem quite like that identified by Maddison and FitzJohn. Here we modify Felsenstein’s original generating process from simple Brownian Motion, to A) Brownian Motion with a single burst occurring on the stem branch of one of the two clades (indicated by vertical dash). B) The distribution of trait values produces a figure very similar to Felsenstein’s original scenario, but results in C) a single contrast (black) that is not well-described by the estimated Brownian Motion process, and thereby generates a significant regression of PIC Y and PIC X (dotted line) despite both X and Y in the shift and BM distributions being uncorrelated. D) As the ratio of the shift variance to the BM variance increases, the proportion of contrast regressions that return a significant result increases dramatically (each point represents 200 simulations for a fixed phylogeny, with both the BM process and the random draw from the shift distribution being uncorrelated with equal variance for both traits). While IC corrects for singular events consistent with Brownian Motion, it does not correct for the more general phenomenon of dramatic singular events driving significant results in comparative analyses. Note that the non-independence of species is not the issue.

**Figure 3:**
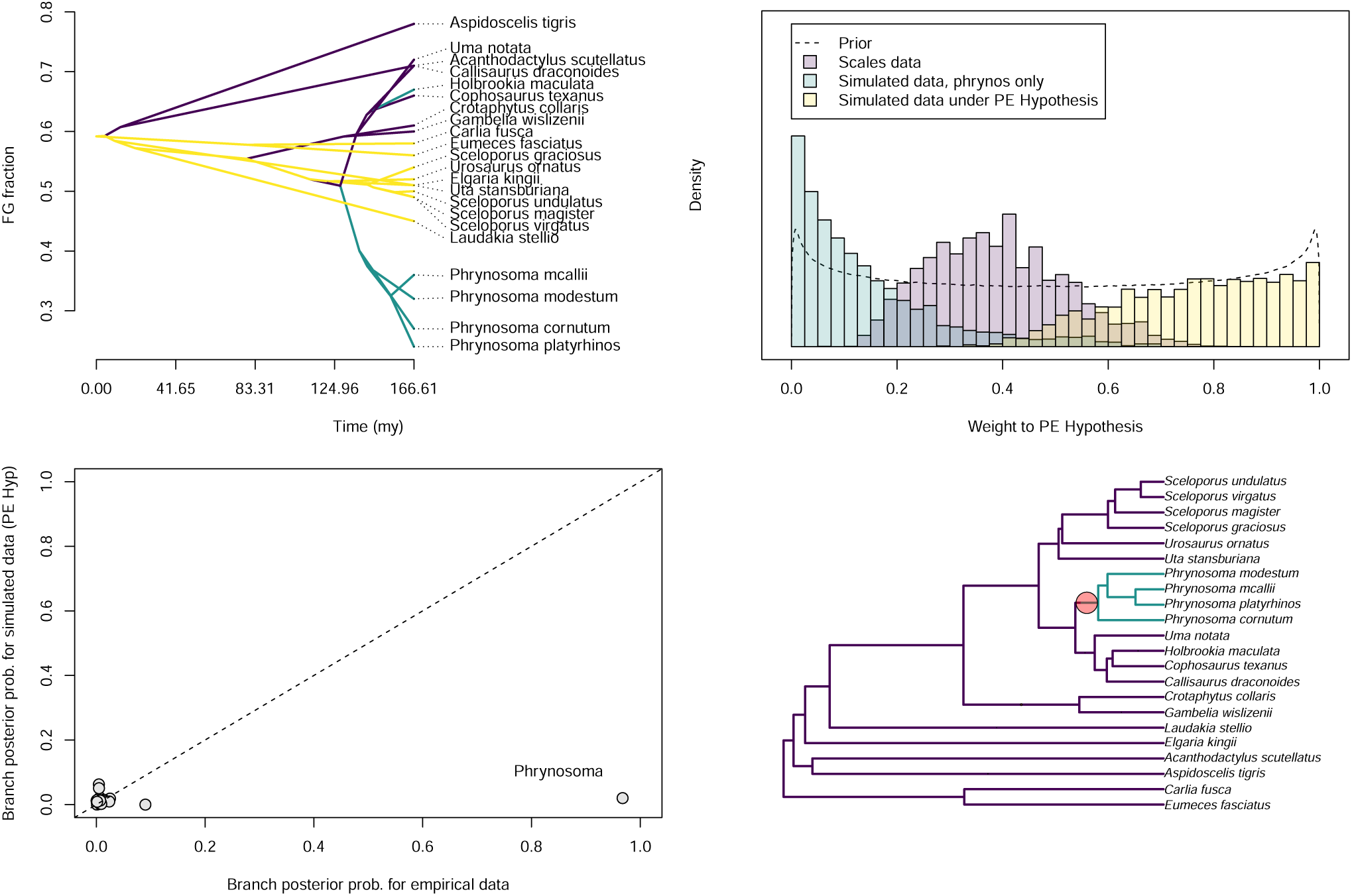
A reanalysis of the Scales et al. (2009) dataset of fast glycolytic muscle fiber fraction across 22 squamate lizards. A) A traitgram depicting the distribution of the data and the reconstructed regimes for the best-fitting Predator-Escape (PE) hypothesis (blue = cryptic, yellow = active flight, purple = mixed). B) Posterior distributions of weights estimated for the PE hypothesis when mixed with a RJMCMC analysis for the original empirical data (purple), data simulated under the best-fitting estimated parameters for a *Phrynosoma*-only shift model (blue), and a dataset simulated under the best-fitting estimated parameters for the full PE model (yellow). Notice that the empirical dataset has intermediate weights. C) Posterior probabilities for all branches of the phylogeny estimated for the original empirical data (X-axis) and the simulated dataset under the PE hypothesis (dashed line is the 1 to 1 line). D) We estimate a high posterior probability on a regime shift in the genus *Phrynosoma* from the empirical data only (red circle), indicating that while the PE hypothesis explains some patterns in the data, it does not fully explain the shift present in the behaviorally and ecologically unique genus *Phrynosoma*.

**Figure 4:**
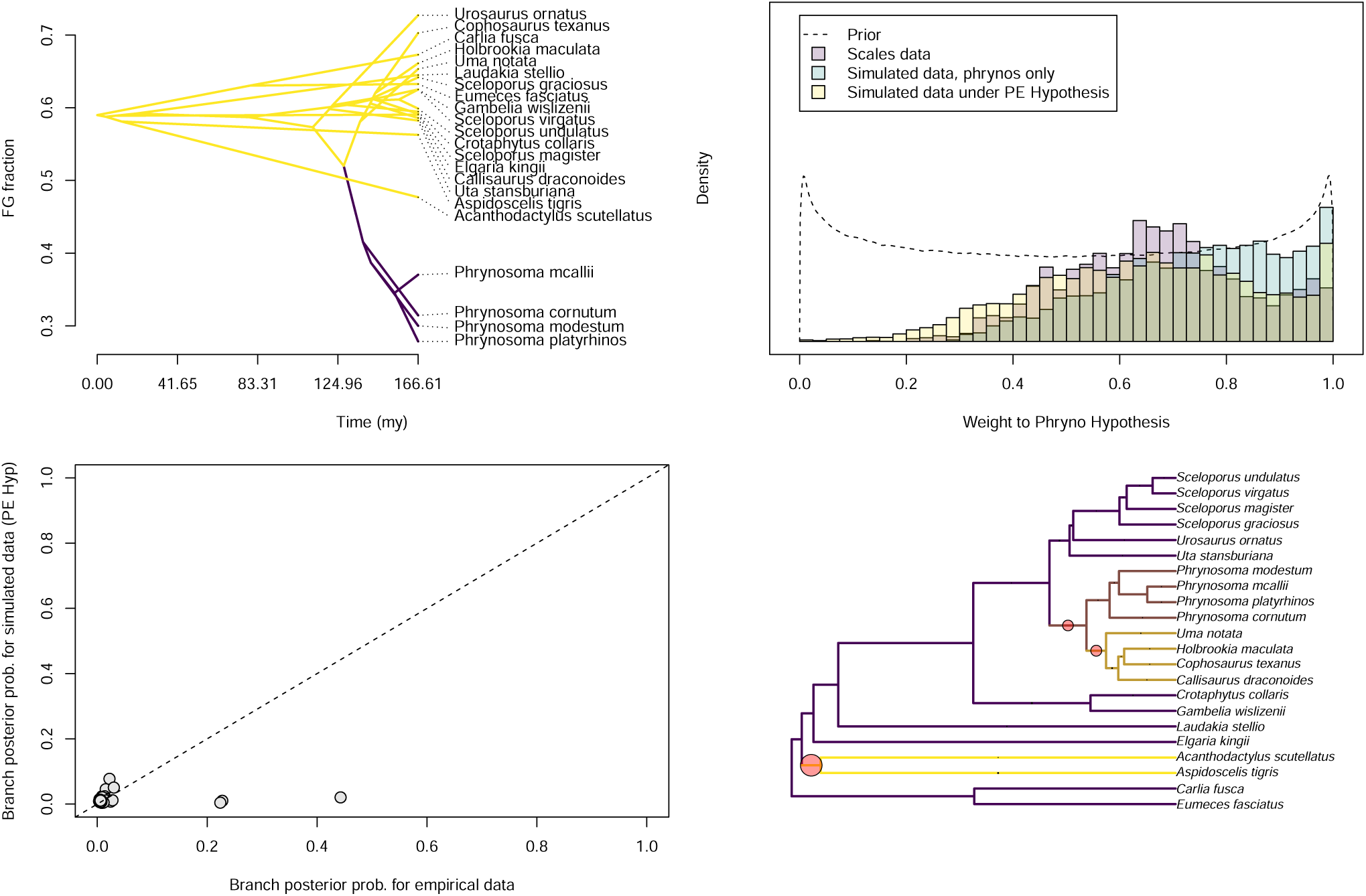
A reanalysis of the Scales et al. (2009) dataset of fast glycolytic muscle fiber fraction across 22 squamate lizards against the *Phyrnosoma*-only hypothesis. A) A traitgram depicting the distribution of simulated data under the *Phyrnosoma*- only hypothesis (yellow = squamates, purple = *Phyrnosoma*). B) Posterior distributions of weights estimated for the *Phyrnosoma*-only hypothesis when mixed with a RJMCMC analysis for the original empirical data (purple), data simulated under the best-fitting estimated parameters for a *Phrynosoma*-only shift model (blue), and a dataset simulated under the best-fitting estimated parameters for the full PE model (yellow). All analysis recover high weights. C) Posterior probabilities for all branches of the phylogeny estimated for the original empirical data (X-axis) and the simulated dataset under the PE hypothesis (dotted line is the 1 to 1 line). D) Modest support for two additional shifts are recovered for the empirical data only (red circles).

**Figure 5:**
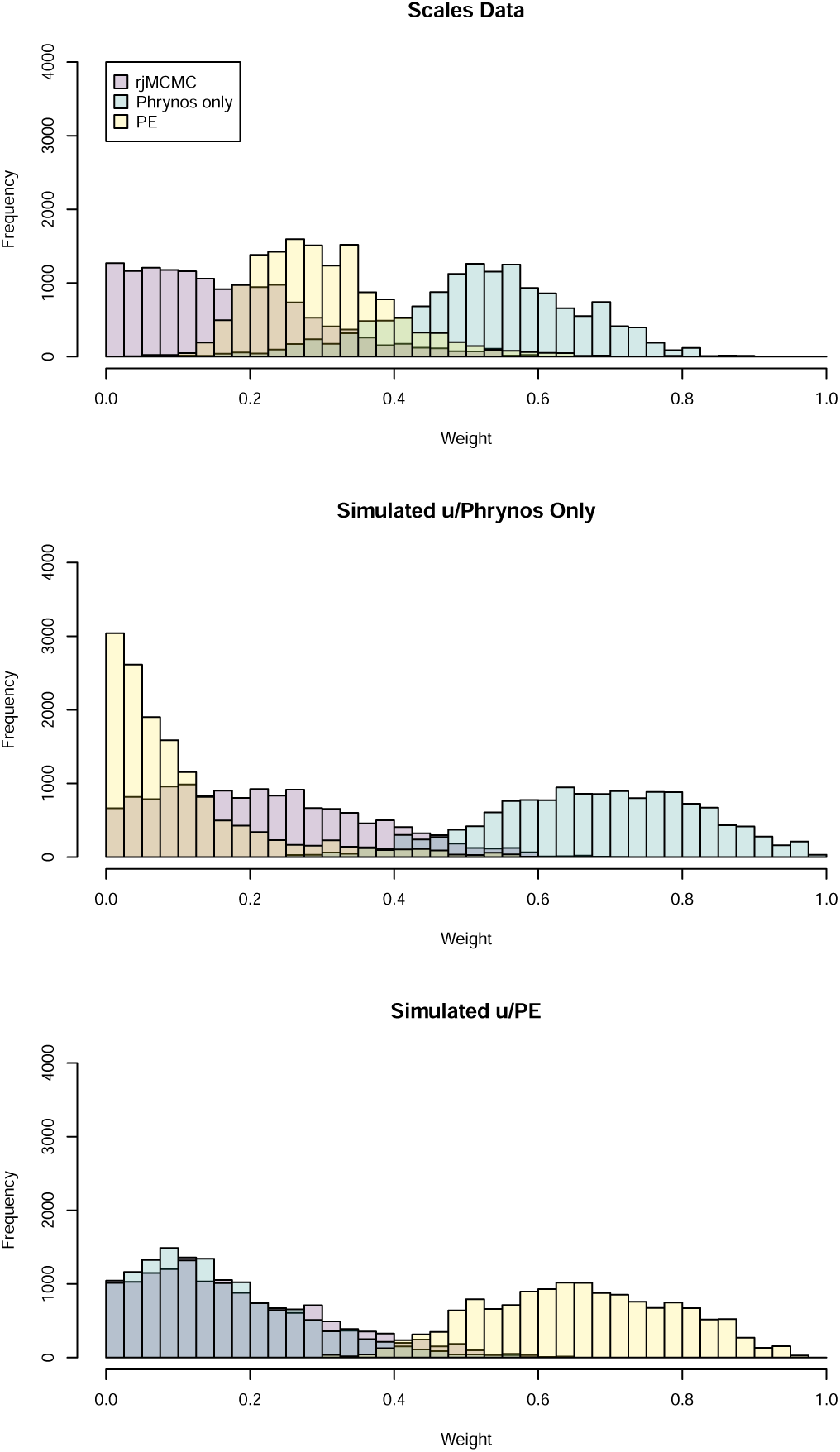
A reanalysis of the Scales et al. (2009) dataset of fast glycolytic muscle fiber fraction across 22 squamate lizards against with both the *Phyrnosoma*-only hypothesis and the PE hypotheses. Weights are depicted for each of the three datasets A) the original Scales dataset B) A dataset simulated under the *Phrynosoma*-only model C) A dataset simulated under the PE hypothesis. In B and C, the correct model receives highest support with neither of the alternatives being well-supported. In the original Scales dataset, the *Phrynosoma*-only hypothesis receives the most weight (indicating a singular shift best explains the patterns observed in the data), while an intermediate weight is given to the PE hypothesis (which explains a good amount of the remaining variation). In no analysis did the reversible-jump portion recover support for any additional shifts.

**Figure 6:**
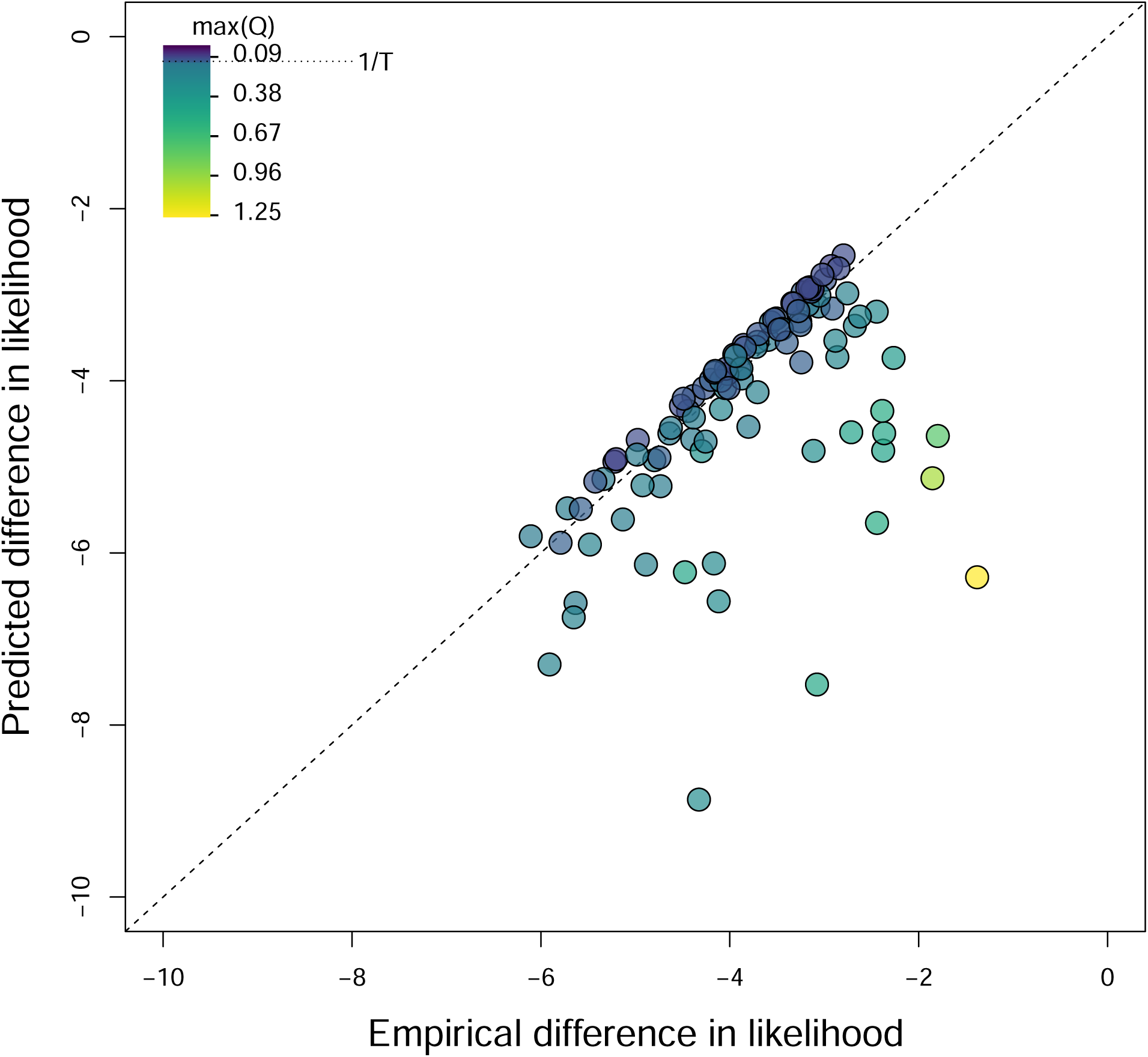
Darwin’s scenario–the singular origin of two coextensive traits on the phylogeny–represents a boundary case to finding the correlation between discrete characters. Pagel’s correlation test for Darwin’s scenario can essentially be reduced to the difference in probability between choosing the same branch twice vs. choosing the branch only once. We demonstrate that here, showing our predicted differences in log likelihood between the independent and dependent trait models (y-axis) against the empirical estimates of the difference in log likelihood between models for simulated Darwin’s scenarios on different phylogenies. Dotted line indicates equality. Points falling off the line represent slight violations of the assumptions we used to derive our prediction. Particularly, we assume that the rates of gain of the traits are so low that only one shift is ever observed. The color of the points indicates cases where this assumption is violated, as outlying points with max(Q) values much greater than 1/*T* (where only 1 shift is expected) are much more likely to fall off the predicted line.

**Figure 7:**
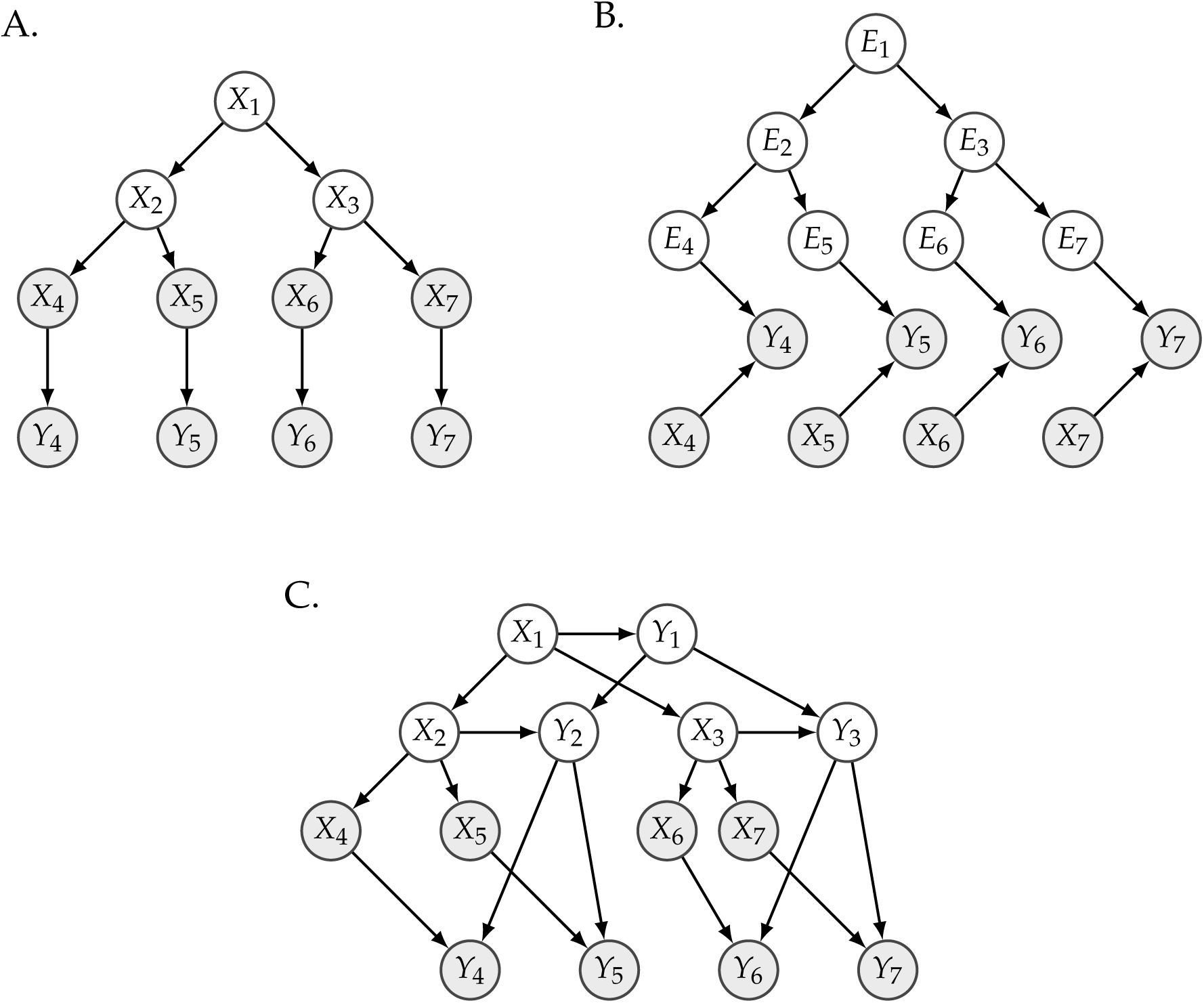
Graphical models of alternative causal relationships between a predictor (X) and a trait of interest (Y). Note that each node has independent, uncorrelated error as an input, but these have not been shown for clarity. A) X follows the phylogeny with observed states (gray) and unobserved ancestral states (white) and is a cause of trait Y. However, the phylogeny and pattern of evolution of X are irrelevant, and this graph can be modeled with methods such as Ordinary Least Squares regression. B) The trait Y has unobserved causes (*E*_*i*_) that follow the phylogeny (gray) that can be modeled using, for example, Brownian motion. The trait X is a cause of Y. This graph can be modeled using methods such as PGLS and PIC. C) The trait Y evolves on the phylogeny and is affected by trait X all throughout its history. Thus, the history of both X and Y must be modeled (e.g. Brownian Motion of X and Ornstein-Uhlenbeck for Y). This graph can be modeled using methods such as SLOUCH.

While other researchers had hit upon similar notions throughout the early 1980s (e.g., Clutton-Brock and Harvey, 1980; Mace et al., 1981; Ridley, 1983; Stearns, 1983; Cheverud et al., 1985), none of these had the pervasive impact that Felsenstein’s presentation did (see for example, Losos, 2011, who reproduces the figures and the accompanying reasoning in his presidential address for the American Society of Naturalists). The problem is just so obvious — data from different clades clustering in different parts of the bivariate plot — all you have to do is look. And while of course his proposed solution, “independent contrasts” (IC), was widely adopted, we suspect it is the clarity with which Felsenstein articulated the problem that has kept his paper a hallmark of biological education and a testament to the importance of tree-thinking, even as his method has largely been superseded by the least squares (Grafen, 1989) (which is identical to IC if BM is used to model the covariance of errors: Rohlf, 2001; Blomberg et al., 2012) and mixed model (Lynch, 1991; Housworth et al., 2004; Hadfield and Nakagawa, 2010) approaches.

However, an important part of this story is often missed: Felsenstein also noted that the problem of non-independence does not occur if “characters respond essentially instantaneously to natural selection in the current environment, so that phylogenetic inertia is essentially absent” (p. 6). Despite this comment, a common misunderstanding of his argument is that the problem inherent in a non-phylogenetic regression of phylogenetically structured data is that species are not independent. In fact, independence of data is not an assumption of standard (non-phylogenetic) linear regression at all. Rather, standard linear regression assumes that the *errors* of the fitted model are independent and identically distributed (i.i.d.). As a result, many applications of a “phylogenetic correction” seem to be missing the point (Revell, 2010; Hansen and Bartoszek, 2012): if all of the phylogenetic signal in a dataset is present in the predictor trait and the errors are i.i.d., then there is no need for any phylogenetic correction (Rohlf, 2001, 2006). (However, phylogenetic analyses are nearly always needed to determine this condition in the first place.)

We suggest that what made Felsenstein’s *prima facie* argument so compelling was that it appealed to biologists’ intuition that many large clades of organisms are just different in many potentially idiosyncratic ways (Vermeij, 2006). If the apparent association between traits found in a non-phylogenetic regression analysis is simply a result of these idiosyncratic differences between clades, then we would be inferring a relationship from unreplicated data (Nee et al., 1996), irrespective of the purely statistical consideration of whether errors are i.i.d.

Here we revisit Felsenstein’s worst-case scenario in order to demonstrate that IC and PGLS do not completely address the problem that we tend to think they do — these methods are still susceptible to singular evolutionary events. To demonstrate this, we add a slight twist to Felsenstein’s original example. First, we used a phylogeny with two clades, each of which is internally unresolved, similar to that of the 1985 paper. We emphasize that the only phylogenetic structure is that stemming from the deepest split. We then simulated two traits under independent BM processes, each with an evolutionary rate (*σ*^2^) of 1. However, at some point on a stem branch of one of the two clades we introduce a singular evolutionary “event” — i.e., a dramatic shift in a lineage’s phenotype — drawn from a multivariate normal distribution with uncorrelated divergences and equal variances that are a scalar multiple of *σ*^2^. The resulting distribution of the data suggests a situation very similar to Felsenstein’s worst-case scenario — and what we suspect is the type of problem envisioned by most biologists when they warn their students of the dangers of ignoring phylogeny.

One would hope that our tools for “correcting for phylogeny” would recognize that the apparently strong relationship between the two traits in our example was driven by only a single contrast. However, this is not the case. That single contrast results in a very high-leverage statistical outlier that drives significance as the size of the shift increases (Figure 2). We can repeat the same exercise with more phylogenetically structured data (where the two clades of interest are fully bifurcating following a Yule process) and obtain identical results (Figure 2, see Supplementary Material). This is disconcerting since our intuition suggests that we do not have compelling evidence for a causal relationship between these two traits (i.e., there is very little reason for us to believe from this correlation alone that one trait is an adaptation to the other).

How can we formulate a better set of models that can account for what our intuition tells us is a dangerous situation for causal inference? We can do so by including another phylogenetically plausible model: that trait correlations result from a single random shift, drawn from a different distribution than the one used to model trait evolution across the rest of the branches.

Let us consider a situation quite distinct from Felsenstein’s multivariate BM (mvBM) scenario. Here traits do not evolve by mvBM, but rather undergo a shift at a single point (perhaps an ancient dispersal event where one clade invaded a new environment). In such a scenario, we only need to consider the phylogeny in as much as a given species exists on either side of the event in question. We can then erect two statistical models: a linear regression model and a singular event model.

Linear regression model:

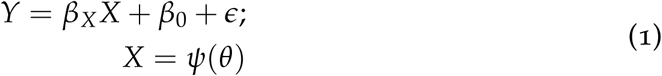

where *β*_*X*_ and *β*_0_ are the slope and intercept to the regression of *Y* on *X, ϵ* is a vector containing i.i.d. random variables that describe the errors, and the predictor *X* is generated by some stochastic process *ψ*(*θ*) on the phylogeny (e.g., a random variable describing a single burst in *X* on the stem branch of one of the two clades with parameters *θ*). Thus, under the laws of conditional probability, the bivariate probability of X and Y under the linear model conditional on parameters *θ*_*LM*_ = (*β*_*X*_, *β*_0_, *σ*) is:

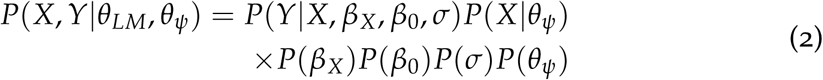

where *θ*_*ψ*_ are the parameters of the process for *X* on the phylogeny, and *σ*^2^ is the variance associated to *ϵ*. This equation is derived from the assumed path of causation between X and Y, since the likelihood function of trait X, denoted by *P*(*X*|*θ*_*ψ*_), is independent of Y, while the likelihood function of Y, denoted by *P*(*Y*|*X, β*_*X*_, *β*_0_, *σ*) depends on X. The remaining terms in the probability statement are interpreted as prior distributions for the parameters in a Bayesian inferential framework.

Alternatively, *X* and *Y* may not be related to one another at all. Rather, they may be the products of singular random evolutionary events denoted by *E*1, and *E*2, that happened to occur on the branch separating two clades.

Singular events model:

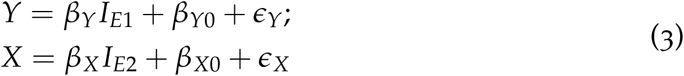

where the variables *I*_*E*1_ and *I*_*E*2_ are indicator random variables that take the value of 1 if an observation is from a lineage that experienced a phylogenetic event or shift, and a value of 0 otherwise. Furthermore, *β*_*Y*0_ and *β*_*X*0_ are the parameters that describe the trait means had they not experienced the singular evolutionary event in question and each linear model in equation (3) has errors with variances 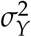 and 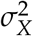 respectively.

For the singular event model (with parameters Λ_*SE*_ = (*β*_*Y*_, *β*_*Y*0_, *β*_*X*_, *β*_*X*0_, *σ*_*Y*_, *σ*_*X*_)) the bivariate probability becomes:

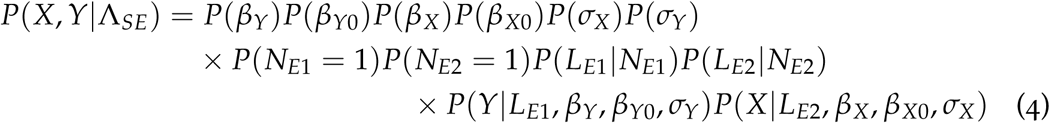

where *P*(*N*_*E*1_ = 1) and *P*(*N*_*E*2_ = 1) are the probabilities of observing a single shift on the phylogeny, and *P*(*L*_*E*1_|*N*_*E*1_) and *P*(*L*_*E*2_|*N*_*E*2_) are the probabilities of observing these singular shifts in locations *L*_*E*1_ and *L*_*E*2_, respectively.

The linear regression and singular event models lead to potentially very different distributions of trait data at the tips. For example, under the the singular event model, the distribution of Y is conditionally independent of X after accounting for *L*_*E*1_, *β*_*Y*_, *β*_*Y*0_ — a testable empirical prediction that will often result in these two models being easily distinguishable with model selection. But failing to consider the singular event model as a possibility is a problem: even for the simple case of two continuous traits, we have shown how easily data simulated under the singular event model can result in highly significant regressions for OLS, PGLS and IC regressions, regardless if the errors are simulated as independent or phylogenetically correlated with respect to the model and phylogeny. We also note that estimating a *λ* transformation for the errors (Pagel, 1999; Freckleton et al., 2002) will not rescue the analysis; the estimated value of *λ* will lie between 0 and 1 and we have found both these more extreme cases (OLS and IC, respectively) to be susceptible.

One might argue that the situation we describe is the violation of the assumption of a BM model of evolution — and this would, of course, be correct (see also Maddison and FitzJohn, 2015). Indeed, for decades it has been common practice (but unfortunately, not universally so) to test whether contrasts are i.i.d. after conducting an analysis using IC (Garland et al., 1992; Purvis and Rambaut, 1995; Slater and Pennell, 2013; Pennell et al., 2015) and many researchers have followed Jones and Purvis (1997) in dropping outlying contrasts from regressions. Felsenstein recognized this particular vulnerability in his method, and correctly predicted that the underlying model was an “obvious point for future development” (p. 14). While today we have a much wider range of comparative models to choose from including some that allow for adaptive shifts, most continuous trait models are Gaussian (e.g., Pagel, 1999; Blomberg et al., 2003; Butler and King, 2004; O’Meara et al., 2006; Eastman et al., 2011; Beaulieu et al., 2012; Uyeda and Harmon, 2014) and do not accommodate abrupt, discontinuous shifts in phenotypes. It is only recently that alternative classes of models have been considered (Landis et al., 2012; Elliot and Mooers, 2014; Schraiber and Landis, 2015; Boucher et al., 2017; Duchen et al., 2017; Blomberg, 2017). Whether or not these other types of models can sufficiently account for rare, singular events will be examined in the next section.

Nevertheless, our primary point here is to suggest that the phenomenon that made Felsenstein’s argument so intuitive is not the violation of i.i.d. errors but rather the biologically intuitive realization that unreplicated differences co-localized on a single branch provide only weak evidence of a causal relationship between traits. Furthermore, models that actually describe such scenarios — like our “singular events” model — are rarely considered in comparative analyses. Admittedly, fitting such models to biologically realistic cases more complex than Felsenstein’s scenario will require estimating the location and number of events and we therefore view our “singular events” model as primarily an illustrative alternative solution to Felsenstein’s thought experiment. Nevertheless, the example illustrates that the phylogeny imposes a challenge to the inference of meaningful associations between traits not because it renders errors non-independent, but because the structure of the phylogeny allows for ancient, potentially unknowable causal factors (which may be few or even singular) to drive widespread associations between traits. Evaluating the validity of these associations as evidence for a meaningful relationship, even in the case of continuous traits, is precisely the unresolved challenge identified by Maddison and FitzJohn (2015) in the case of discrete character correlations (as we will further elaborate in Case Study III).

### Case Study II: Adaptive hypotheses and singular shifts

As stated above, the IC method is based on the BM model of trait evolution. While this model is useful (and has often been used) for testing for adaptation, it is inconsistent with how we think of the *process of adapting* to an optimal state (Lande, 1976; Hansen, 1997; Hansen and Orzack, 2005; Hansen et al., 2008; Hansen and Bartoszek, 2012). Hansen’s introduction of the Ornstein-Uhlenbeck (OU) process to comparative biology and the suite of methods built on his approach have been the only real attempts to actually try and capture the basic dynamics of adaptive trait evolution on phylogenies.

Multi-optima OU models have been widely used to test for the presence of shifts in evolutionary regimes (i.e., parts of the phylogeny with their own optima, or less commonly, their own strength of selection parameters). Tests of adaptive evolution come in two flavors: those with an *a priori* hypothesis (or hypotheses) regarding which lineages belong to which distinct regimes based on ancestral state reconstruction of explanatory factors (Butler and King, 2004; Beaulieu et al., 2012) and those where the locations of regime changes are themselves estimated along with the parameters of the OU process (Ingram and Mahler, 2013; Uyeda and Harmon, 2014; Khabbazian et al., 2016).

These two types of approaches represent two different philosophies of data analysis that follow a schism that cuts through comparative methods. For example, there are two major ways to investigate the dynamics of lineage diversification: test specific hypotheses about the drivers of diversification rate shifts (for example, the ‘SSE’ family of models; Maddison et al., 2007; FitzJohn, 2012) or search for the most-supported number and configuration of shifts (Alfaro et al., 2009; Stadler, 2011; Rabosky, 2014). The former (hypothesis-testing) seeks to understand the causes of evolutionary shifts, while the latter (data driven) is a descriptive and exploratory approach to understanding evolutionary patterns. As we alluded to above, we refer to these data-driven approaches as “phylogenetic natural history” due to their similarity to the practice of natural history observations in nature but projected backwards through phylogenetic space and time (Maddison and FitzJohn, 2015).

Of course, the types of inferences we can make will be limited by our choice of approach. For example, it may be tempting to use exploratory approaches such as *BAMM* (Rabosky, 2014) or *bayou* (Uyeda and Harmon, 2014) to search a vast range of model space to find a particularly well-supported statistical hypothesis, observe the shifts identified, and then come up with post hoc explanations for why that particular configuration fits an adaptive story that the researcher can suddenly construct with great precision. (Comparative biologists are of course not unique in succumbing to such temptations; see for example Pavlidis et al., 2012). In fact, discovering the location of well-supported shifts on the phylogeny does not say anything about causation; it is merely a descriptive technique to find major features of the data where there is evidence that the parameters governing the dynamics of trait evolution have shifted on the phylogeny. It is nonetheless useful — and we argue essential — that a researcher know where these shifts occur. The reasons for this are covered in Case Study I: these major shifts are likely to drown out any biological signal in a dataset if they are unaccounted for by our hypothesis-driven models. But even beyond these statistical considerations, observing which lineages and clades differ in evolutionary tempo and mode is as vital to good macroevolutionary inference as traditional natural history is to biology more generally — such knowledge and familiarity with the organisms is essential to generating “empirically-justifiable” synthetic theories of evolution (Futuyma, 1998). For these reasons, we argue that hypothesis-driven and phylogenetic natural history approaches are complementary: we must pit our particular causal hypotheses against a descriptive “stuff-happens” model built on idiosyncratic singular evolutionary events.

To illustrate how we might go about uniting these two modes of inference to disentangle the support for causal models of evolution from that attributable to singular events, we reanalyze a dataset introduced by Scales et al. (2009) on lizard muscle fiber proportions (hereafter, the ‘Scales’ dataset). (An expanded dataset was re-analyzed by Scales and Butler (2016) with slightly modified hypotheses. However the original 2009 paper serves as a clearer example with which to illustrate our perspective; we do not delve into differences between the two.)

Scales et al. (2009) are interested in the composition of muscle fiber types in squamate lizards, and whether these muscle fibers evolve adaptively in response to the changing behavior and ecology of the organisms. They propose three primary adaptive hypotheses for the drivers of fast glycolytic (FG) muscle fiber proportions: i) foraging mode behavior (FM; e.g., sit-and-wait vs. active foraging vs. mixed); ii) predator escape behavior (PE; e.g., active flight vs. crypsis vs. mixed); and iii) a combined hypothesis of foraging mode and predator escape (FMPE) that assigns a unique regime to every combination of FM and PE represented in the dataset. For each hypothesis, they reconstruct a likely phylogenetic history of these behavioral modes on the phylogeny by conducting ancestral state reconstructions (Figure 3). After fitting the multi-optimum OU models to the muscle fiber data, they find strong support for the predator escape hypothesis, which is 13.0 AICc units better than the next closest model (FMPE). Such a finding appears quite reasonable under the “Life-Dinner Principle” (Dawkins and Krebs, 1979), which suggests that escaping a predator may have a far more direct effect on fitness than obtaining a food item (Scales et al., 2009).

However, AIC provides only relative support for a model given a set of alternatives (see Pennell et al., 2015, for more on this point in the context of comparative methods). An examination of the particular configuration of shifts in the three hypotheses may give pause to researchers familiar with squamates. For example, some may want to quibble with the suggestion that the “sit-and-wait” foraging behavior of *Phrynosoma* species, which are often ant-eating specialists that leisurely lap up passing insects, should be grouped with the “sit-and-wait” tactics of species such as *Gambelia wislizenii*, a voracious carnivore that frequently subdues and consumes other lizards close to their own size. Looking at the reconstructions, it is also apparent that the PE hypothesis is the simplest model that allows a shift on the branch leading to *Phrynosoma*, a group that any herpetologist would identify as “weird” for a multitude of reasons (indeed, these are the eyeball-socket-blood-squirters alluded to in the introduction). The question then arises: is the signal in the dataset for the PE hypothesis driven entirely by the singular evolution of different muscle fiber composition in *Phrynosoma* lizards? If so, then any number of causal factors that differ between *Phrynosoma* and other lizards could be equally as likely as predator escape — including foraging mode with a slight reclassification of character states! We want to emphasize that we are not criticizing any of the particular choices the researchers involved in this study made. Rather, we argue that such quandaries are the inexorable result whenever the primary signal in the data is due to a singular historical event.

To explore the impact of the distinctiveness of simply being a *Phrynosoma* lizard, we developed a novel Bayesian model by building on the R package *bayou* (Uyeda and Harmon, 2014). To do so, we consider the macroevolutionary optimum of a particular species to be a weighted average of past regimes, as is typical in all OU models with discrete shifts in regimes (Butler and King, 2004; Beaulieu et al., 2012), but in our case, this weighted average is itself a weighted average of two differing configurations of the locations of adaptive shifts (often referred to as “regime paintings”). One configuration assumes that shifts in the optima have occurred where a discrete character, hypothesized to shape the evolutionary dynamics of the continuous character, is reconstructed to have shifted. The other configuration is estimated directly from the data using *bayou*’s reversible-jump MCMC (RJMCMC) algorithm.

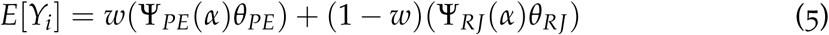

This equation describes the expected value of a trait for species *i, Y*_*i*_ as a weighted average between the expected trait value under the PE hypothesis and the expected trait value under the reversible-jump estimate of regime shift configurations. The vectors *θ*_*PE*_ and *θ*_*RJ*_ are the values of the trait optima for the *N*_*PE*_ and *N*_*RJ*_ adaptive regimes, while Ψ_*PE*_ and Ψ_*RJ*_ correspond to the standard OU weight matrices that average over the history of adaptive regimes experienced by species *i* over the course of their evolution, with older regimes being discounted proportional to the OU parameter *α* (for a full description of how these weight matrices are derived, see Hansen, 1997; Butler and King, 2004).

In our model, the regime painting for our a priori hypothesis Ψ_*PE*_ is fixed, while we estimate the parameters the configuration of shifts for the reversible-jump component, Ψ_*RJ*_, as well as the values for the optima *θ*_*PE*_ and *θ*_*RJ*_; and standard parameters for the OU model such as *α* and *σ*^2^ which are assumed constant across the phylogeny. We also estimate the weight parameter *w*, which determines the degree of support for the PE hypothesis against the reversible-jump regime painting. We place a truncated Poisson prior on the number of shifts for the reversible-jump analysis to be quite low, with a mean of *λ* = 0.5 and a maximum of *λ* = 10 (meaning that we are placing a prior expectation of 0.5 shifts on the tree). Furthermore, we place a symmetric *β*-distributed prior on the *w* parameter with shape parameters of (0.8, 0.8). Additional details on the model-fitting can be found in the Supplementary material.

We then fit this model to 3 different datasets: i) the original Scales data; ii) data simulated using the Maximum Likelihood estimates for the parameters of the PE model fitted to the Scales dataset; and iii) data simulated under the Maximum Likelihood estimates for a “*Phrynosoma*-only” model in which a single shift occurs leading to the genus *Phrynosoma*. We then compared the posterior distribution of the weight parameter *w* to evaluate the weight of evidence for each hypothesis in each dataset.

We find that our approach places intermediate weight on the PE hypothesis for the original Scales dataset. When we simulated data under the PE hypothesis, the estimated weight given to the PE hypothesis was likewise high (Figure 3B). When data were simulated under the *Phrynosoma*-only hypothesis, the weight given to the PE hypothesis was low, as predicted (Figure 3B). Furthermore, the RJ portion of the model fit to the Scales dataset recovers only a single highly supported shift on the stem branch of the *Phrynosoma* lizards (Figure 3C and 3D). This suggests that the PE hypothesis has statistically supported explanatory power as its estimated weight is well bounded away from 0. But it does not explain everything. In particular, the PE hypothesis fails to fully explain the shift leading to the *Phrynosoma* lizards (Figure 3C and 3D), which are more extreme than they should be considering the other taxa in their regime (there is only one, *Holbrookia maculata*, which does not show such an extreme shift). In summary, the signal for an association between muscle fiber composition and predation escape behavior is generated in part, but not completely, by variation that is specific to the genus *Phrynosoma*. Therefore, the conclusions of Scales et al. (2009) hold up, even when accommodating evolutionary regime shifts unrelated to the factors being considered, although the weight of evidence appears to be larger than it actually is. This more subtle view of muscle fiber evolution conforms quite well with our biological intuition — variation in predator escape behavior is a good explanation for observed patterns of muscle fiber divergence, but *Phrynosoma* are a unique group with other factors likely influencing their trait evolution beyond predator escape.

We can conduct the same analysis where we test not the PE hypothesis, but the *Phrynosoma*-only hypothesis against the reversible-jump hypotheses (Figure 4). In this case, we recover high weights for the *Phrynosoma*-only hypothesis regardless if the model is fit to the Scales dataset, or to data simulated under either the *Phrynosoma*-only hypothesis or the PE hypothesis. This is because accounting for the *Phrynosoma* shift is the primary feature of all three datasets (though weights are somewhat higher for data simulated under the *Phrynosoma*-only hypothesis than others). It may appear unsatisfying that such high weights are recovered for the a priori hypothesis when a singular event, which is easily reconstructed by the RJMCMC, explains the distribution of the data just as well.

However, the analysis favors the *Phrynosoma*-only hypothesis simply because of the vague priors placed on the number and location of shifts in the reversible-jump analysis. Guessing correctly which of the 42 branches on the phylogeny has a single shift with our hypothesis is rewarded by the analysis (we will return to this issue in Case Study III). In the original Scales dataset, there are weakly supported shifts in the clades leading to the sister group of *Phrynosoma* lizards, and the branch leading to *Acanthodactylus scutellatus* and *Aspidoscelis tigris*. Finally, we can combine all three hypothesis simultaneously by placing a Dirichlet prior on the vector *w* = [*w*_*RJ*_, *w*_*PE*_, *w*_*Phrynosoma*_]. Doing so recovers strongest support for the *Phrynosoma*-only model, intermediate support for the PE hypothesis, and very little weight on the reversible-jump hypothesis, which has no strongly supported shifts (Figure 5).

By combining phylogenetic natural history approaches with our a priori hypotheses, we show that we can account for singular evolutionary events that are not well-accounted for by our generating model. In the case of the PE hypothesis, we show that it does indeed have explanatory power beyond simply explaining a singular shift in *Phrynosoma* and support the original authors’ conclusions. However, the intermediate result likely only occurs because the PE hypothesis places *Phrynosoma* in the same regime as *Holbrookia maculata*, which does not share the extreme shift that is found in *Phrynosoma*. Were this not the case (as in our fitting of the *Phrynosoma*-only hypothesis), it would still require visual inspection of the phylogenetic distribution of traits under the hypothesis in question to determine that a singular evolutionary event is driving support for a particular model. As discussed above, given a large enough tree such a priori hypotheses are likely to be strongly supported; if you can predict which one branch out of many will contain a shift then you may be on to something. But given the dangers of ascertainment bias and our biological intuition, we find this interpretation unsatisfying(Maddison and FitzJohn, 2015). As with Case Study I, these scenarios ultimately reduce to whether or not coincident unreplicated events are evidence of a causal link between traits (Maddison and FitzJohn, 2015), a problem we will address in Case Study III.

Nevertheless, we show the value in combining a hypothesis testing framework with a natural history approach to identifying patterns of evolution. We show here that allowing for unaccounted shifts can provide a stronger test and more nuanced conclusions regarding the support for a particular predictor driving trait evolution across a phylogeny. Furthermore, predictors which provide additional explanatory power (if for example, regimes are convergent or if predictors vary continuously) will be even more favored over natural history models. Thus, our framework certainly does not automatically reward more complex, freely estimated models. Rather, the great uncertainty in possible models is incorporated as a prior on the arrangement of shifts and is limited in explanatory power, some-thing that researcher-driven biological hypotheses are much more capable of accomplishing.

### Case Study III: Darwin’s scenario and unreplicated bursts

We now turn to a case where both the explanatory variable and the focal trait are discrete characters. Detecting a signal of evolutionary covariation is more difficult in discrete characters, but examining this situation isolates the recurring problem we have identified in the two previous Case Studies — whether or not coincident unreplicated events are evidence of causal links between traits. As we mention above, Maddison and FitzJohn (2015) recently demonstrated that commonly used methods return significant correlations all the time — and in scenarios that seem to defy our statistical intuition. For example, Pagel’s (1994) correlation test would find the phylogenetic co-distribution of milk production and middle ear bones highly statistically significant even though they both are a defining characteristic of mammals, an inference so obviously dubious that even Darwin (1872) warned against it. This seems to be a clear case of phylogenetic pseudoreplication (Maddison and FitzJohn, 2015; Read and Nee, 1995). More broadly, Maddison and FitzJohn describe the goal of correlation tests as finding the “weak” conclusion that “the two variables of interest appear to be part of the same adaptive/functional network, causally linked either directly, or indirectly through other variables” (p. 128). They assert that with our current approaches, we cannot even clear this (arguably low) bar. Here we delve into this idea a bit deeper. What constitutes good evidence of such a relationship and why precisely do phylogenetic tests for correlations provide such apparently unreasonable results?

Maddison and FitzJohn highlight two hypothetical situations, that they refer to as “Darwin’s scenario” and an “unreplicated burst”. They argue that these scenarios provide little evidence for an adaptive/functional relationship between two traits because the patterns of co-distribution only reflect singular evolutionary events (Figure 1). In Darwin’s scenario, two traits are coextensive on the phylogeny, meaning that in every lineage where one trait is in the derived character state, the other trait is as well. As an example, consider the aforementioned phylogenetic distribution of middle ear bones and milk production in animals; all mammals (and only mammals) have middle ear bones and produce milk. These traits (depending on how they are defined) have only appeared once on the tree of life and both occurred on the same branch (the stem branch of mammals). The unreplicated burst scenario is identical to Darwin’s scenario except that rather than a single transition occurring in both traits, there is a single transition in the state of one trait (e.g., the gain of middle ear bones) and a sudden shift in the transition *rates* in another trait (e.g., the rates by which external testes are gained and lost across mammals). Note that these scenarios do not differ qualitatively from Felsenstein’s worst-case scenario nor the *Phrynosoma*-only model scenario from Case Studies I and II (Figure 1). In all three scenarios, something novel and interesting happened on a single branch and the distribution of traits at the tips of the phylogeny reflects this.

In their paper, Maddison and FitzJohn (2015) simulated comparative data and reported a preponderance of significant results using Pagel’s correlation test (1994) and Maddison’s (1990) concentrated changes test. In order to hone our intuition of the problems they present, we dig a bit deeper and investigate the mathematical reason that Pagel’s discrete correlation test (1994) returns a significant result in Darwin’s scenario. (We should note here that Brookfield [1993] conducted a similar analysis that was more-or-less completely overlooked.) To make the problem tractable, we assume that the traits were selected for study without first looking at their phylogenetic distribution, a condition that we (as well as Maddison and FitzJohn, 2015) suspect is rarely met in practice (more on this below).

Again, under Darwin’s scenario, there is a single concurrent origin of two traits leading to perfect co-distribution across the phylogeny for all taxa stemming from branch (a condition we define mathematically as event *A*). Under the independent model, both traits *X* and *Y* have to switch from 0 to 1 on the same branch *L* once. For these traits, we can make the assumption that the likelihood of evolving these traits at all is quite small. In other words, replaying the tape of life, under Markovian assumptions, will likely lead to many worlds where milk and middle ear bones don’t exist at all. However, we do not study traits that don’t exist. Thus, for traits such as these, we can expect that there is likely to be only one origin of the trait on the phylogeny. Thus, the probability of the independent model denoted as*P*(*M*_*ind*_|*A*) given the assumptions above is

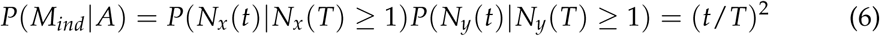

where *t* is the branch length of branch *L* containing both shifts (Karlin and Taylor, 1981) and *N*_*x*_ and *N*_*y*_ are the stochastic processes that denote the number of shifts of trait *X* and *Y* at time *t* respectively (see Supplementary Material for exact derivation of the probability the independent model). Since this is the only way Darwin’s scenario can occur on a branch for *M*_*ind*_, the probability in Eq. (6) is equivalent to the likelihood value *L*(*M*_*ind*_) evaluated at the maximum likelihood estimate of *M*_*ind*_.

In contrast, for the completely dependent model *M*_*dep*_, it is enough to follow what happens in a single trait since the second will just simply change along. The probability of the dependent model *P*(*M*_*dep*_|*A*) under Darwin’s scenario is

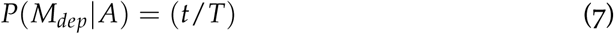

(see full derivation of this probability in the Supplementary Material). The likelihood value at the maximum likelihood estimate of this model turns out to be just the probability from Eq. (7), that is *L*(*M*_*dep*_) = *t*/*T*. Therefore, the test statistic *D* used in the likelihood ratio test for Pagel’s discrete correlation test comparing models *M*_*ind*_ and *M*_*dep*_ is simply proportional to the ratio of the length of the branch where the shift occurred to the total length of the tree:

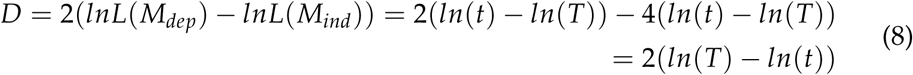

In other words, the results of the analysis are predetermined. Under Darwin’s scenario, including additional taxa in the analysis will increase the support for the dependent model simply as a consequence of increasing the total length of the tree (i.e., because *T* increases and the difference between *ln*(*T*) and *ln*(*t*) will get larger as long as the additional sampled taxa do not break Darwin’s scenario).

The assumptions used to derive this result differ very slightly from those used in available software; however, we can use simulation to test the validity of our result and to demonstrate that this is the mathematical reason that Pagel’s test returns a significant result. Using the R package *diversitree* (FitzJohn, 2012), we simulated a set of 20 taxon trees where both traits underwent a irreversible transition on a single, randomly chosen, internal branch. We then fit a Pagel model with constrained (*M*_*dep*_) and unconstrained (*M*_*ind*_) transition rates. We also constrained the root state in both traits to 0, rates of losses of both the traits to 0, and gain rates in the dependent model following the gain of the other trait to be extremely high. Plotting the empirically estimated differences in the MLEs against the predictions making the simplifying assumptions above reveals a strong modal correlation between them (Fig. 6). Differences likely reflect the fact that we have not explicitly made the assumption that *P*(*N*_*x*_(*t*) = 1) = *P*(*N*_*y*_(*t*) = 1) ≈ 1 when we fit the model with *diversitree*. Furthermore, we compare here only fully dependent and independent models. This can be seen when calculating the probability of one switch in each trait *P*(*N*_*x*_(*t*) = 1, *N*(*t*)_*y*_ = 1). In the fully dependent case that simply becomes *P*(*N*_*x*_(*t*) = 1), in the independent case it becomes *P*(*N*_*x*_(*t*) = 1)*P*(*N*_*y*_(*t*) = 1) but in the correlated case it becomes *P*(*N*_*y*_(*t*) = 1 *N*_*x*_(*t*) = 1)*P*(*N*(*t*)_*x*_ = 1) ≠ 1 affecting the likelihood ratio test based on estimations of the correlation (see Supplementary Material). However, such intermediate cases will only introduce slight differences and may not be distinguishable from the fully dependent case under Darwin’s Scenario (though they will be important in more intermediate cases, see Supplementary Material).

Maddison and FitzJohn (2015) hinted that the coincident occurrence of single events could be a way of measuring the evidence for a correlation, but did not work out the details as we have done here. The key to understanding the above results is to recall Gould and Eldredge’s famous dictum (1977) that “stasis is data”. The remarkable coincidence is not just that the two characters happened to evolve on the same branch but that they were never subsequently gained or lost through-out the rest of the tree. For even a modestly sized tree, this coincidence is so unlikely that the alternative hypothesis of correlated evolution is preferred over the null. It is therefore not completely unreasonable that Pagel’s test tells us that these traits have evolved in an entirely correlated fashion.

However, one key consideration should make us suspect of this line of reasoning. As Maddison and FitzJohn (2015) point out, the traits we use in comparative analyses are not chosen independently with respect to their phylogenetic distribution (as we assumed in our analysis). Rather, researchers’ prior ideas about how traits map unto trees likely inform which traits they choose to test for correlated evolution. For example, it is common practice among systematists to search for defining and diagnostic characteristics for named clades; these traits are of especial interest and are likely the same sorts of traits that are researchers might include in comparative analysis, thereby greatly increasing the likelihood of finding traits with independent, unrelated origins that align with Darwin’s scenario. We agree with Maddison and FitzJohn (2015) that this type of ascertainment bias is likely prevalent in empirical studies, even if it is usually more subtle than testing for a correlation between milk and middle ear bones. However, the presence of ascertainment bias does not mean it is not worth attempting to discover the source of the signal. Understanding the exact mathematical reasons why Pagel’s test infers a significant correlation in a given case provides a clear boundary condition that can help develop quantitative corrections for ascertainment bias. Furthermore, the issues of ascertainment bias are likely to rapidly dissipate as we move away from the boundary case of Darwin’s scenario. As a result, extending our analytical approach to more complicated scenarios will likely provide an even more meaningful estimate of the weight of evidence supporting a hypothesis of correlation.

#### The structure of a solution

We have shown in the three Case Studies that many PCMs, including those that form the bedrock of our field, are susceptible to being misled by singular evolutionary events. This fundamental problem has sown doubts about the suitability and reliability of many methods in comparative biology (e.g., Losos, 2011), even if it was not obvious that these issues were connected. But again, the fact that apparently different issues share a common root makes us hopeful that there can be a common solution.

As we illustrate through our Case Studies, we think that accounting for the possibility of idiosyncratic evolutionary events will be an essential step towards such a solution. However, we will need to think hard about how best to model such events. In Case Study II, we present one solution to the problem that involves explicitly accounting for the possibility of unaccounted adaptive shifts using Bayesian Mixture modeling. We believe this approach has a great deal of promise as it provides simultaneous identification of biologically interesting shifts and the explanatory power of a particular hypothesis.

However, we do not claim that such an approach is the only solution or that it solves the problem completely. Indeed, we find that in all three Case Studies, the uniting philosophy is to consider models that account for background shifts in evolutionary regime, rather than strict adherence to a particular methodology. For example, we highlighted in the introduction that we think HSMs (following Beaulieu et al., 2013; Beaulieu and O’Meara, 2016) are a potentially powerful, and widely applicable solution, even though we did not consider these in detail here.

And there are still other potential solutions which we have not even mentioned yet. In our own work (Uyeda et al., 2017), we have used a strategy similar to the Bayesian Mixture Modeling presented in Case Study II, but instead of modeling the trait dynamics as a joint function of our hypothesized factors and background changes (represented by the RJMCMC component), we did the analyses in a two-step process: first, we used *bayou* (Uyeda and Harmon, 2014) to locate shifts points on the phylogeny, then used Bayes Factors to determine if predictors could “explain away” shifts found through exploratory analyses. For PGLS and other linear modeling approaches, modeling the errors using fat-tailed distributions (Landis et al., 2012; Blomberg et al., 2012; Elliot and Mooers, 2014; Duchen et al., 2017) may mitigate the impact of singular evolutionary events on the estimation of the slope (also see Slater and Pennell, 2013, for an alternative approach using robust regression). Furthermore, we also think that rigorous examination of goodness- of-fit and model adequacy following any comparative analysis is critical for finding unforeseen singular events driving signal in the dataset (Garland et al., 1992; Boettiger et al., 2012; Slater and Pennell, 2013; Pennell et al., 2015). Which of these solutions (including those that were included in our Case Studies and those that were not) will be the most profitable to pursue will probably differ depending on the question, dataset and application — we anticipate that there will not be a one-size-fits-all solution — but we do think that any compelling solution will involve a unification of phylogenetic natural history and hypothesis testing approaches.

But we want to take this a step further. While it is useful to account for phylogenetic events in our statistical models, a greater goal of comparative biology should be explain why these events exist in the first place. We return to Maddison and FitzJohn’s (2015) “weak” goal of finding whether or not “two variables of interest appear to be part of the same adaptive/functional network, causally linked either directly, or indirectly through other variables.” We ultimately disagree with them that this constitutes a weak conclusion; the challenges of making these inferences from any comparative dataset are significant. Furthermore, we find the often repeated axiom “correlation does not mean causation” to be un-helpful. While it is accurate in the strict sense, some patterns of correlations will *at least be consistent* with a given model of causation whereas others will not be. And it is clear from reading the macroevolutionary literature, biologists do not shy away from forming causal statements from correlative data regardless. While some might reasonably wish us to simply highlight these claims as violating statistical principles, we think it is worthwhile to take seriously the question: “What would it take to infer causation from comparative data?” And even if we are to conclude that all the evidence for a hypothesized causal relationship stems from one or a few evolutionary events, is this finding biologically meaningful?

#### Phylogenies are graphical models of causation

One way to gain a foothold on the problem of causation is to build, communicate, and analyze phylogenetic comparative methods in a graphical modeling frame-work — a perspective that has recently been advocated by Höhna et al. (2014, 2016). Graphical models that depict hypothesized causal links between variables make explicit key underlying assumptions that may otherwise remain obscured; indeed, the precise assumptions of PCMs were hotly debated in the early days of their development (Westoby et al., 1995b,a; Nee et al., 1996; Harvey et al., 1995; McNab, 1988; Westoby, 2007) and remain poorly understood to this day (Hansen and Orzack, 2005; Hansen and Bartoszek, 2012). As examples of how using graphical models force us to be more clear in our reasoning, consider the graphs in Figure 7. We depict three different models of causation that have phylogenetic effects that each require alternative methods of analysis to estimate the effect of trait X on trait Y. In our example, a four species phylogeny provides possible pathways for causal effects, but variables may have entirely non-phylogenetic causes or may be blocked from ancestral causes by observed measurements, rendering the phylogeny irrelevant (e.g. Figure 7A). Edges connect nodes and indicate the direction of causality, where the nature of phylogenies allows us to assume that ancestors are causes of descendants, and not vice versa. This asymmetry results in a what is known as a probabilistic Bayesian Network (a type of directed acyclic graph, or DAG) that predicts a specific set of conditional probabilities among the data.

Depending on the Bayesian network structure, the appropriate method of analysis can range from a non-phylogenetic regression (Figure 7A), to commonly used comparative methods such as Phylogenetic Generalized Least Squares (PGLS, Figure 7B), to methods that require modeling both the evolutionary history of interaction of both trait X and trait Y (Figure 7C) (Hansen, 1997; Butler and King, 2004; Hansen et al., 2008; Revell, 2010; Hansen and Bartoszek, 2012). We emphasize that this implies that the use of phylogeny in interspecific comparisons is an *assumption* that depends on the precise question being asked and the hypothesized causal network. It is often assumed and asserted that PCMs are simply a more rigorous version of standard regression. This is simply not true.

In cases where phylogeny does matter, we must specify the generating model for unobserved states in our causal graphs. For example, it is common to assume a BM model for residual variation in PGLS or that ancestral states are reconstructed using stochastic character mapping in OU modeling of adaptation. However, BM and other continuous Gaussian or Markov processes are only a few of the many types of processes that may generate change on a phylogeny. We have shown that discontinuous processes and singular events are poorly handled in our current framework and lead to much confusion about what exactly, our statistical methods are allowing us to infer from comparative data. Such models can be similarly illustrated using graphical models (Figure 8). By making our models explicit, we see that the phylogeny is best thought of as a pathway for past factors to causally influence the present-day distribution of observed states. These “singular-event” models are alternatives to the more continuous models we typically examine. Furthermore, representing our models as graphs, we are poised to take advantage of the sophisticated approaches for causal reasoning (e.g., Pearl, 1995, 2009; Sugihara et al., 2012; Shipley, 2016) that subsume familiar tools such as path analysis, structural equation modeling, and graphical modeling into a more generally applicable structural theory of causation. These tools have been embraced by fields such as computer science, epidemiology and the social sciences, but largely ignored by comparative biologists (rare exceptions are the recent introductions of phylogenetic path analysis by Hardenberg and Gonzalez-Voyer [2013] and the application of causal models to make inferences from paleontological time-series by Reitan and Liow [2017]).

**Figure 8:**
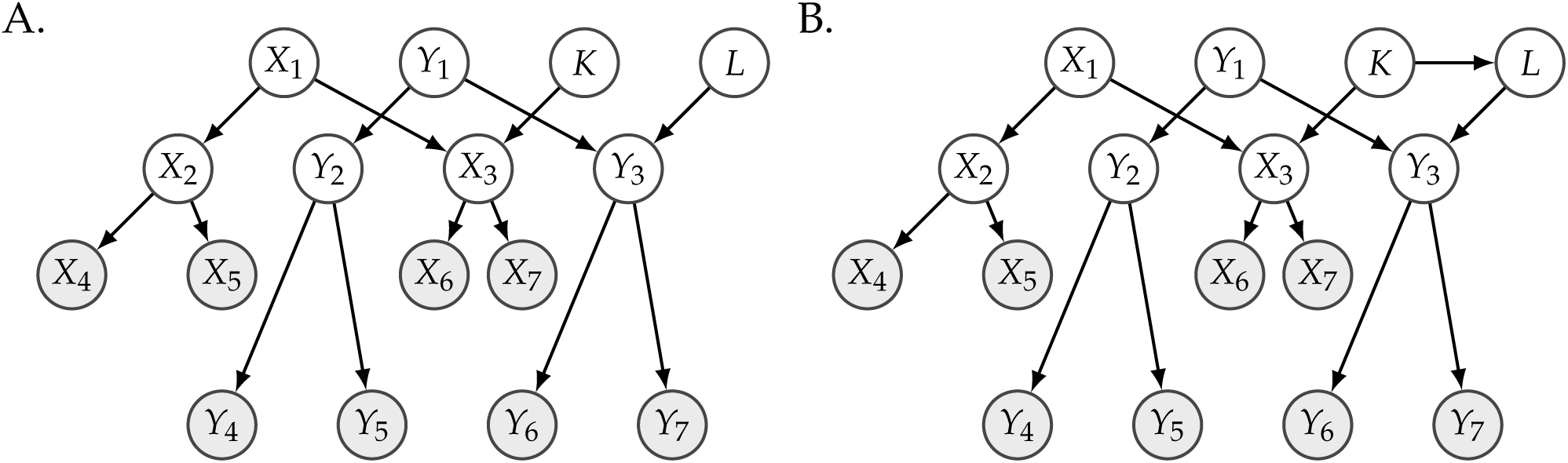
Graphical models of Darwin’s Scenario between a predictor (X) and a trait of interest (Y). Note that each node has independent, uncorrelated error as an input, but these have not been shown for clarity. A) Singular event model. Here two independent factors cause a change on ancestral states *X*_3_ and *Y*_3_ (*K* and *L* respectively). However, they are independent events and coincidentally occur at the same point on the phylogeny. B) Similar to the previous model, but *K* and *L* are causally linked. Thus, whenever *K* occurs, it probabilistically causes *L* which causes a shift in *Y*. If only one event occurs however, this model is only distinguishable from graph (D) proportional to the probability that events *K* and *L* occur on the same branch (see Case Study III).

One clear case where such graphical modeling would improve inference are cases where considering phylogeny reverses the sign of the relationship between two variables. This is precisely what Nee et al. (1991) found looking at the relationship between body size and abundance in British birds; depending on how they aggregated the data (means of species, means of genera, means of tribes, etc.) the direction of correlation flipped back and forth. This reversal in the sign of the relationship between two variables *X* and *Y* when conditioning on a third *Z* is a general, and widely studied, statistical phenomenon known as “Simpson’s paradox” (Blyth, 1972). Nee and colleagues (1991; 1996) hold up their findings of the British bird study to be emblematic; in their view, the presence of Simpson’s paradox in their data clearly implies that phylogeny is key to making sense of interspecific data.

However, as Pearl (2014) has convincingly demonstrated, Simpson’s paradox is not really paradoxical at all when considered from the standpoint of Bayesian Networks. In fact, Pearl shows that the appropriate way to analyze the data depends crucially on what one assumes is causing what. To understand how causal inference resolves Simpson’s Paradox, we now present a rather artificial, but nevertheless illustrative example (Pearl, 2009). Consider three traits: Body size (*B*), abundance (*N*) and migratory behavior (*M*) in birds. Given the Bayesian Networks presented in Figure 9, we have two possible hypotheses for the causal relationships between the traits. We further consider the possibility that we do not have adequate data on *M*, and thus only *B* and *N* are observed. Our goal is to estimate the causal effect of *B* on *N*. In Figure 9A, body size influences whether or not species become migratory, and both migratory status and body size influence species abundance (but in opposite directions). Furthermore, under this scenario, both body size and migratory status will have phylogenetic signal. We can evolve traits along the phylogeny depicted in Figure 9C and obtain a bivariate plot that looks like Figure 9D. Under the alternative Bayesian Network, migratory behavior still has a positive effect on species abundance, but also increases body size, which in turn causes decreases species abundance. These two causal structures are observationally equivalent — meaning that any distribution simulated under one can be replicated under the alternative causal structure. Therefore, both networks can produce datasets with phylogenetic signal in both body size and migratory behavior, and both can produce a dataset with the distribution in Figure 9D (see Supplementary Material for additional details on generating Figure 9).

**Figure 9:**
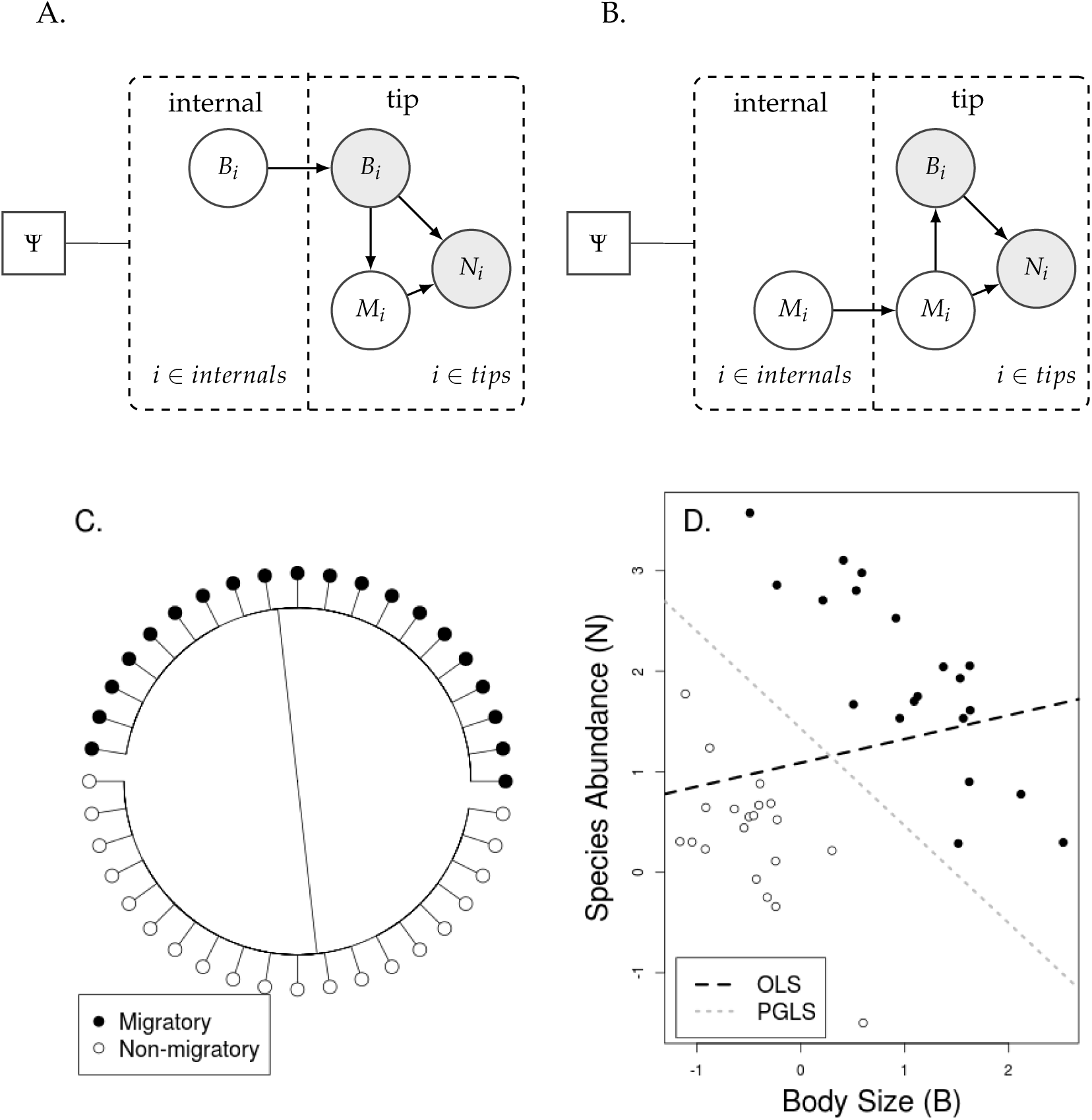
Simpson’s paradox in phylogenetic comparative methods. Panels (A) and (B) depict two alternative Bayesian Networks. In (A), body size is a cause of both species abundance and migratory behavior, and trait B evolves on the phylogeny Ψ. We represent evolution on the phylogeny using a “tree plate” (see Höhna et al., 2014, dashed box) where (unobserved) node states can influence tip states. In (B), body size still affects species abundance, but migratory behavior itself is a cause of both body size and species abundance, but the phylogenetic effect is present in migratory behavior (in this case, we simulated with a Brownian threshold model). (C) A phylogeny similar to that of Darwin’s scenario used to simulate the dataset (D), with migratory species (black) and non-migratory (white) taxa. The data in (D) can be generated by either causal structure. However, to estimate the effect of B in both networks, one must use different analytical approaches. To estimate the net effect of B on N in network (A), the appropriate method of analysis is OLS regression (black line). This is because increasing body size will simultaneously decrease species abundance and increase migratory behavior, which itself increases abundance, leading to a net slight increase in abundance. However, under network (B) the correct method is PGLS (gray line) as increasing body size will have no effect on migratory behavior, and unaccounted phylogenetic residual error is present in the observed data. Here, increasing body size will only have a direct effect of decreasing species abundance, which is reflected in the estimate of the slope. The resolution of Simpson’s paradox rests entirely on causal assumptions; which are immediately apparent from graphical models but difficult to express with standard mathematical formulae.

How then should we analyze the data if we want to understand the effect of body size on species abundance? If we assume that body size influences migratory behavior, then increasing body size (for example, if natural selection leads a species to become larger) will increase the probability of that species becoming migratory — and the two opposing effects will result in relatively little change in species abundance. Therefore, we should perform Ordinary Least Squares regression to estimate the net causal effect of increasing body size. We also note that all the phylogenetic signal is coming from the evolution of body size, which becomes irrelevant once we observe body size, and thus we do not need to perform PGLS. By contrast, if migratory behavior causes changes in body size, then selecting for an increase in body size will not result in a lineage changing their migratory status at all. Therefore, we are assured that increasing body size will likewise always decrease species abundance. Consequently, we should perform PGLS to account for the phylogenetic signal in the residual variation imposed by (unobserved) migratory status.

By working through the logic of comparative analyses using graphical models we have come to essentially the same line of reasoning of Westoby et al. (1995b,a), who, in the early days of PCMs, challenged the growing consensus that phylogeny needed to be included in any interspecific comparison — a consensus which has only gotten stronger as the years passed by (also see McNab, 2003, for a related critique). Westoby and colleagues were concerned that including phylogeny in interspecific comparisons necessarily favored some causal explanations over others. At the time, their critique was dismissed as innumerate hogwash (Harvey et al., 1995; Nee et al., 1996) and this evaluation has largely stuck. However, from our example of bird size and abundance, it is apparent that Westoby et al. were right all along: phylogenetic comparative methods are a powerful tools for drawing inferences from interspecific data but they necessarily imply some types of causal structures and negate others. It is too much to ask of our methods to decide what questions we ought to ask. As Westoby et al. (1995a) put it: “No statistical procedure can substitute for thinking about alternative evolutionary scenarios and their plausibility” (p. 534).

#### Concluding remarks: are our models valid tests of our causal hypotheses?

By explicitly including phylogeny into our graphical models of causation, we are forced to reckon with the scope of the inference problem and the ability of our data to be informative. While most of the statistical assumptions of methods are often well-known (e.g., for linear models, we assume that errors have equal variance and are normally distributed, etc.), Gelman and Hill (2006) argue that there is a more fundamental assumption — validity of data — that is almost always implicit and often overlooked:

> “Most importantly, the data you are analyzing should map to the research question you are trying to answer. This sounds obvious but is often overlooked or ignored because it can be inconvenient. Optimally, this means that the outcome measure should accurately reflect the phenomenon of interest, the model should include all relevant predictors, and the model should generalize to the cases to which it will be applied.” (Gelman and Hill, 2006)

We believe that far less discussion in comparative methods has been focused on the issue of statistical validity of the data collected to the research questions being posed by a given study. This is in large part because comparative data and the phylogeny that underly it are largely beyond the control of the researcher, but careful consideration of the data is required to understand what research questions can be reasonably answered. For example, we find that most comparative research questions have a poorly defined scope of inference: it is unclear to what population a model or inference should generalize to. If we ask “are milk and middle ear bones correlated?”, we must also specify “in what organisms?”. Since no organisms other than mammals have the particular traits we define as “milk” and “middle ear bones”, we actually do not need statistics at all to determine whether these traits are correlated — we have sampled nearly the entire population relevant to the question! In nature, they are perfectly collinear. If we wish to expand our scope of inference to hypothetical organisms that evolve milk and/or middle-ear bones we are free to do so. However, we have collected a very poor data sample for such a question. It is not the fault of the statistical method to demonstrate that a poorly designed experiment does not represent its scope of inference, rather it is our job as researchers and statisticians to ask whether or not such a relationship addresses our biological question and whether the sample of data collected is valid for the question being asked.

In this paper we have tried to synthesize a wide variety of statistical and philosophical concepts to lay out a roadmap for where we think comparative biology should go. We certainly do not have all the answers. Of the paths we have explored, there are many details that need to be worked out, and we fully anticipate that there are many alternative paths that we have not even considered. However, we argue that if we are going to make substantial progress in using phylogenetic data to test evolutionary hypotheses, we will need to reckon more seriously with the idiosyncratic nature of evolutionary history, and to more clearly articulate precisely what we want to test and whether our models and data are suitable for the task.

## Code Availability

Data and code needed to reproduce all analyses in this manuscript are available at https://github.com/uyedaj/pnh-ms/.

## Acknowledgments

We thank Luke Harmon, Dolph Schluter, Mark Westoby, Daniel Caetano, Eliot Miller, Ben Freeman, Florent Mazel, Joel McGlothlin, Martha Muñoz, Barbara Neto-Bradley, Francisco Henao Diaz, Mauro Sugawara, Emma Goldberg, Michael Landis, Simon Blomberg, and Nicholas Matzke for their critical feedback on these ideas and this manuscript. JCU would like to specially thank the insightful knowledge and teaching gleaned from conversations over the years with Thomas Hansen that inspired the bulk of this manuscript (though he holds no culpability for the contents and opinions therein). MWP was supported by a NSERC Discovery Grant. JCU was supported by NSF Grants to Luke Harmon (DEB-1208912) and JCU (DBI-1661516). RZF was supported by NSF Grant to Luke Harmon (DEB-1208912).

## Supplementary Methods

### A. Case Study I-Supplementary Methods

In order to construct phylogenies that correspond to “Felsenstein’s worst-case scenario”, we simulated two phylogenies with a Yule process and a birth rate of 1 for 20 species and transformed them using Pagel’s *λ* values of either 0 (polytomies) or 1 (fully bifurcating). Trees were then scaled to unit height and combined into a single, two-clade phylogeny with stem branches of equal length (again, unit height). The entire phylogeny was then scaled to unit height so that each clade begins diversifying after 0.5 units of tree height. We ran 200 simulations for each tree type and shift value (either full polytomies or fully bifurcating) where we simulated a bivariate Brownian Motion process with two uncorrelated traits with *σ*^2^ = 1 for both traits. We then chose one of the two stem branches and simulated a shift. We tested 10 different values of the shift variance in an increasing sequence such that each value is 10 times larger than the last, ranging from *σ*^2^ = 10^*-*2^ to 10^3^, from an uncorrelated bivariate Normal distribution. Thus, for each combination of shift variance (10 values) and phylogeny (2 trees) we ran 200 simulations. For each dataset, we then performed Phylogenetic Independent Contrasts (PICs) and estimated the P-value for the slope of the linear regression, forcing the intercept through the origin.

### B. Case Study II-Supplementary Methods

We analyzed the dataset of Scales et al. (2009) using the trait *FG*.*frac* and matched to the squamate phylogeny of Pyron and Burbrink (2014). While this is a different phylogeny than was used in (Scales et al., 2009), the topology was identical and branch length differences were minimal. We implemented a novel approach for combining hypothesis-testing and exploratory reversible-jump MCMC in the software package *bayou*. To do so, we developed a customized R code (available at https://github.com/uyedaj/pnh-ms).

Priors on parameters are as follows: *α* ∼ half-Cauchy(scale = 0.1); *σ*^2^ half- ∼ Cauchy(scale = 0.1); *k* truncated Poisson(*λ* = 0.5, *K*_*max*_ = 10); *θ* ∼ Normal(*µ* = 0.5, *σ* = 0.25); *w* ∼ Beta(shape1 = 0.8, shape2 = 0.8). Each branch of the phylogeny was given an equal probability of a shift with a uniform probability on a given branch (i.e. shifts were allowed to occur anywhere on a given branch with equal probability).

In addition to the empirical dataset, we fit each model to two simulated datasets. First, we simulated under the a *Phrynosoma*-only model that contained a single shift at the base of the stem branch leading to the genus *Phrynosoma*. We chose parameter values close to the values estimated when fitting a *Phrynosoma*-only model model to the original Scales dataset. Specifically, we set *α* = 0.15, *σ*^2^ = 0.001, *θ*_*root*_ = 0.6, and *θ*_*Phrynosoma*_ = 0.3. Second, we simulated a dataset under the PE hypothesis with the same parameters of *α* and *σ*^2^, but with *θ*_*mixed*_ = 0.5, *θ* _*f light*_ = 0.7 and *θ*_*cryptic*_ = 0.3. Again, these values are very close to the estimates we obtained from fitting the PE model to the original Scales dataset.

We ran 3 analyses in each of the 3 datasets, resulting in 9 total MCMC chains. First, we ran the PE hypothesis against the reversible-jump analysis for all 3 datasets. Next we ran the *Phrynosoma*-only hypothesis against the reversible-jump analysis for all datasets. Then we ran a model that places a Dirichlet prior on the weights of all three: PE, *Phrynosoma*-only and RJMCMC. This prior was set to *w* ∼ *Dirichlet*(0.33, 0.33, 0.33) with all other priors being identical to the other analyses. We ran each MCMC chain for at least 200,000 generations or until adequate effective sample sizes were obtained for all parameters (>100) and inspected each chain for evidence of poor mixing. Given the small size of the dataset and very few number of RJMCMC shifts, MCMC chains tended to converge quickly.

### C. Case Study III-Supplementary Methods and Results

As described in the main text, we used a number of simplifying assumptions to generate the prediction that the difference in likelihoods between the dependent and independent cases in Darwin’s scenario is a simple function of the length of branch *L* and the total length of the tree, *T*. To demonstrate this effect, we simulated Darwin’s scenario and performed constrained likelihood optimizations in an effort to come as close as possible to the generating assumptions we made in deriving our result (Figure 6). To do so, we simulated 100 phylogenies under a Yule process with birth rate = 1. We then scaled the tree to unit height and randomly selected an interior branch at which both traits *X* and *Y* were gained. We then estimated the likelihoods for *M*_*ind*_ and *M*_*dep*_ as described in the main text by taking the Maximum Likelihood of 5 replicated optimizations using the *optim* method in R. These replicates were performed to help minimize optimization errors.

As described in the main text, the estimates in Figure 6 were obtained by constraining the models to be either completely dependent or completely independent, for rates of gains between traits to be equal, and by constraining losses of traits to 0 (irreversible). These assumptions were made based on the argument that under Darwin’s scenario there is no evidence from the data that would reject these assumptions. To verify that this is indeed the case, we present the results of an unconstrained model that does not impose these restrictions. Here, the only constraint on the model is that we specify that the root state of the model is state “00” (absence of both traits). Thus, under this model the independent case has 4 transition rates and the dependent case has 8 transition rates (compared to only 1 and 2 in the analysis in the main text, respectively). We used *nlminb* to maximize the likelihoods of these functions, as the increased number of parameters resulted in very flat likelihood surfaces and slow optimization—necessitating the use of bounds (all transition rates were bounded between 0 and 1000). Analyzing the same simulations as before, we obtain a very similar pattern Figure S1, although more cases of the dependent model have higher likelihoods than the independent model (as evident by more points falling above the predicted 1-to-1 line). We conclude that the simplifying assumptions we used to obtain our result in Case Study III are sound, and that the primary reason the dependent model is favored over the independent model is the difference in placing one event on branch *L* and the probability of placing two events on branch *L*.

#### Probabilities of observing one transition in a single branch under independent and dependent models

For a single trait we have that the infinitesimal probability matrix for trait *X* is simply defined as

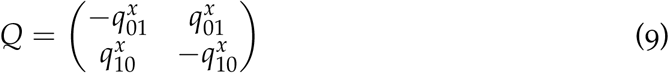

Therefore, the probabilities of the continuous-time Markov chain of trait *X* changing over time in a branch *L* with time *t* are defined via *P*(*t*) = *e*^*Qt*^. In fact, the full probabilities in the irreversible case (when 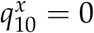) result in transition probability matrix

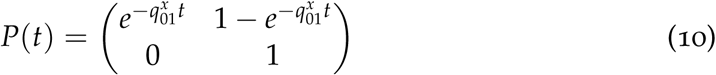

Notice that once branch *L* has switched to state 1 then the probability of the trait staying in state 1 is also 1 (absorbent state). If we want to calculate the number of switches from 0 to 1 trait *X* has experienced along the phylogeny we can define a new stochastic process *N*_*x*_(·) that follows the number of transitions from 0 to 1 through time. This process has a binomial distribution that depends on the number of branches of the tree *B* but also the probability of transition from 0 to 1 in the branch length *t*. That is,

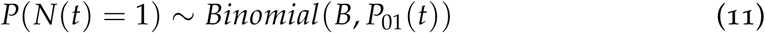

In the case of rare traits the probability of observing a single event in branch *L* is small 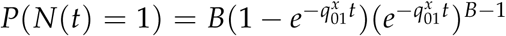 when 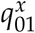 has a small value. However, the probability of observing at least one transition across all the phylogeny, that we denote as the event *N*_*x*_(*t*) ≥ 1 in the main manuscript is simply one minus the probability of zero transitions occurring. Thus,

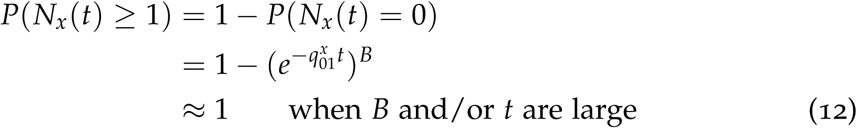

Even when the probability of a switch is small, in the whole tree the probability of observing at least one switch in *X* is almost 1 as demonstrated above (and since we don’t study traits that don’t exist, ascertainment bias will assure that this will be the case for rare traits). The binomial distribution in (Eq. 11) converges to a Poisson distribution with parameter 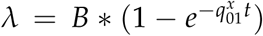 when 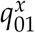 is small and the number of branches *B* is large. If the probability of having more than one transition is negligible but we know there has been at least one transition from 0 to 1, then we are only interested in the location of that single transition. Because the process *N*_*x*_(·) can be also defined via a Poisson process with parameter *λ* as defined above, we know that the probability of one event occurring in an specific branch *L* is simply a uniform distribution based on the length of that branch. Therefore *P*(*N*_*x*_(*t*)|*N*_*x*_(*T*) ≥1) = *t*/*T* where *T* is the sum of all branch lengths. Thus, if the Maximum Likelihood estimate of *λ* is near or below 1, then the probability of observing a single event is nearly 1, and the likelihood of that event is simply the uniform probability of placing that event on branch *L* (the full derivation of the uniform distribution for arrivals in a Poisson processes can be found in Karlin and Taylor (1981)).

Thus, for two traits distributed under Darwin’s scenario (event *A*) shown in Eq. (6), we can calculate the probability of *M*_*ind*_:

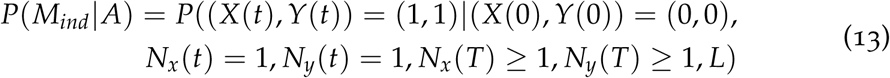

where *N*_*x*_ and *N*_*y*_ are the stochastic processes that denote the number of shifts of trait *X* and *Y* at time *t* respectively, and *L* is the branch on which both transitions occur and it a has a branch length of *t*. Since *X* and *Y* are independent, the joint probability of *X* and *Y* changing at the same time is simply the product of probabilities of each event, therefore the probabilities pertaining *X* and *Y* can be multiplied

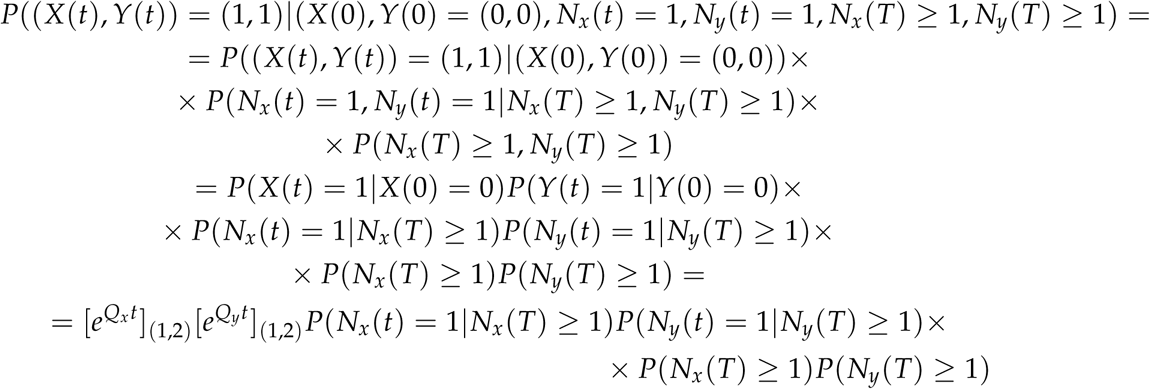

where *Q*_*x*_ and *Q*_*y*_ are the infinitesimal probability matrices that describe the transition rates between states in the independent case as defined in Eq. (10) and the subscripts on 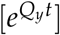(1,2) indicate row 1, column 2 of the resulting probability matrix. We now consider the outcome of maximizing this expression under a likelihood framework. Since there is no evidence of a transition from 1 to 0 in either trait, the maximum Likelihood estimate (MLE) for the transition rates 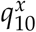 and 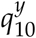 will be 0. Meanwhile, the MLEs 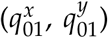 for the transitions from 0 to 1 in both traits will be small (because these events are so rare, occurring only once) but positive since one transition does occur on *L*. Given the resulting parameter estimates 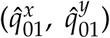, it is likely that a great many realizations of this process would likely result in no lineages evolving the traits of interest at all. However, we do not study traits that don’t exist. (Lewis [2001] followed a similar line of reasoning when he used the Mk model for phylogenetic inference.) Because of this ascertainment bias, the probability of at least one switch occurring for traits that are unlikely to evolve at all (i.e. with very small 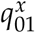 and 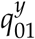) should be nearly exactly one, that is *P*(*N*_*x*_(*t*) ≥1) ≈ 1 when accounting for total branch length *T* of the tree as shown above. The probability of exactly one transition of each trait occurring in the lineage *L* given that at least there is one transition in the tree is simply uniform *P*(*N*_*x*_(*t*)|*N*_*x*_(*T*)≥1) = *t*/*T* as shown above. Furthermore, with rare events the estimates of the probabilities of both traits changing only once in lineage *L* conditional upon observing Darwin’s scenario (under the independent model *M*_*ind*_) is also one 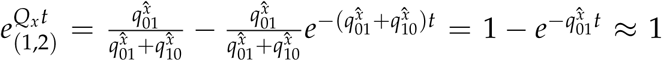 and 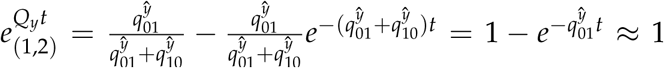, meaning that at the end the probability of the independent evaluated at the maximum likelihood estimate under independent model reduces to

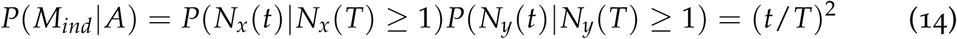

where *t* is the branch length of branch *L* containing both shifts (Karlin and Taylor, 1981).

For the dependent model (*M*_*dep*_), we let the time *t*_*X*_ be the moment at which *X* occurred and the time *t*_*Y*_ be the time at which *Y* occurred in the branch *L* (0 ≤ *t*_*X*_, *t*_*Y*_ ≤ *t*). In Case Study III we can assume that 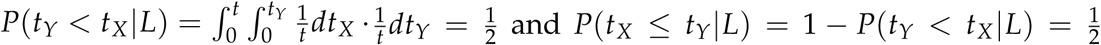. The probability of *M*_*dep*_ under Darwin’s scenario when *X* arrives before *Y* is then:

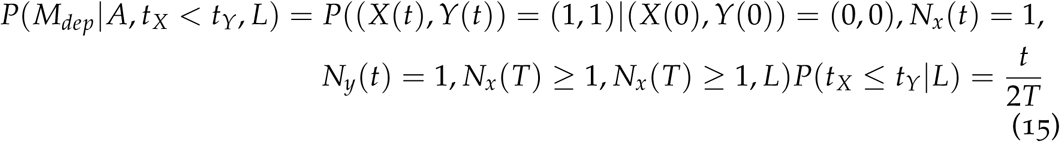

Analogously 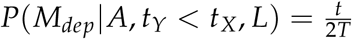 when *Y* happens before *X*. Therefore

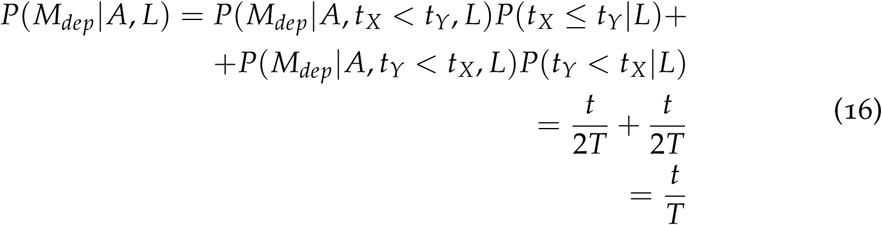

That is the result we denote in Eq. (7). In the main manuscript we omitted writing the dependency on branch *L* for simplification purposes. However, the dependency of a given branch length is discussed throughout Case Study III.

#### Probabilities of observing one transition in a single branch under correlated model

In the correlated full model for Pagel is described via an infinitesimal probability matrix *Q* with eight parameters representing the possible transitions of traits *X* and *Y* that have states 0 or 1.

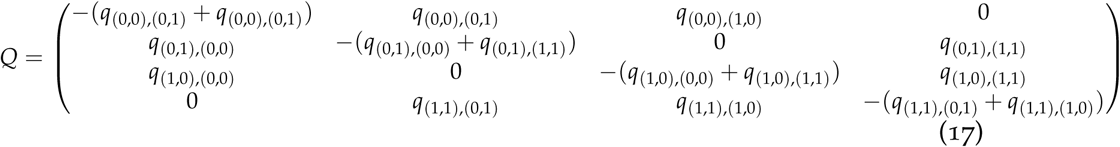

In the irreversible case we have that *q*(0,1),(0,0) = *q*(1,0),(0,0) = *q*(1,1),(0,1) = *q*(1,1),(1,0) = 0, and the *Q*-matrix from (17) is reduced to four parameters. We are interested in calculating the probability of both traits moving to state 1. Therefore

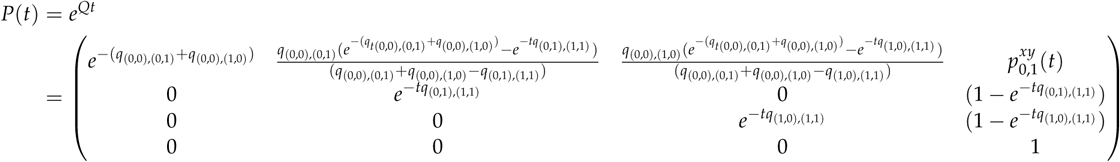

So the probability that we are interested in is

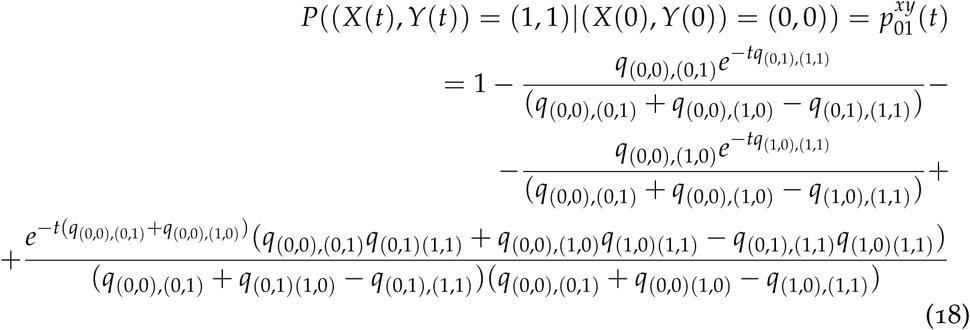

When the branch length *t* is sufficiently large we have that 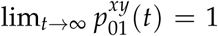 just as the independent and fully dependent cases because *e-tα* → 0 for any *α* > 0.

### D. Graphical Models-Supplementary Methods

To obtain the results in Figure 9 in the main text, we considered two Bayesian Networks involving 3 traits. Body size (*B*) and species abundance (*N*) are observed continuous traits, while a third trait, migratory behavior (*M*) is a threshold trait—meaning that it is a discrete trait that has an underlying continuous liability (Felsenstein, 2011). For the network in Figure 9A, we generated data by simulating Brownian Motion (root = 0, *σ*^2^ = 1) of *B* on the phylogeny depicted in Figure 9C. We then simulated the liability of *M* as *M*_*liab*_ = *B* + *ϵ* where *ϵ* is a random Normal deviate with mean 0 and standard deviation of 0.5. We then discretized *M*_*liab*_ into a binary character (*M*) by assigning liabilities above the median value of *M*_*liab*_ 1 and below the median value 0. Finally, we simulated values of *N* as *N* = *B* + − 3*M* + *ϵ*, where *ϵ* is again a random Normal deviate with mean 0 and standard deviation of 0.5.

**Figure S1:**
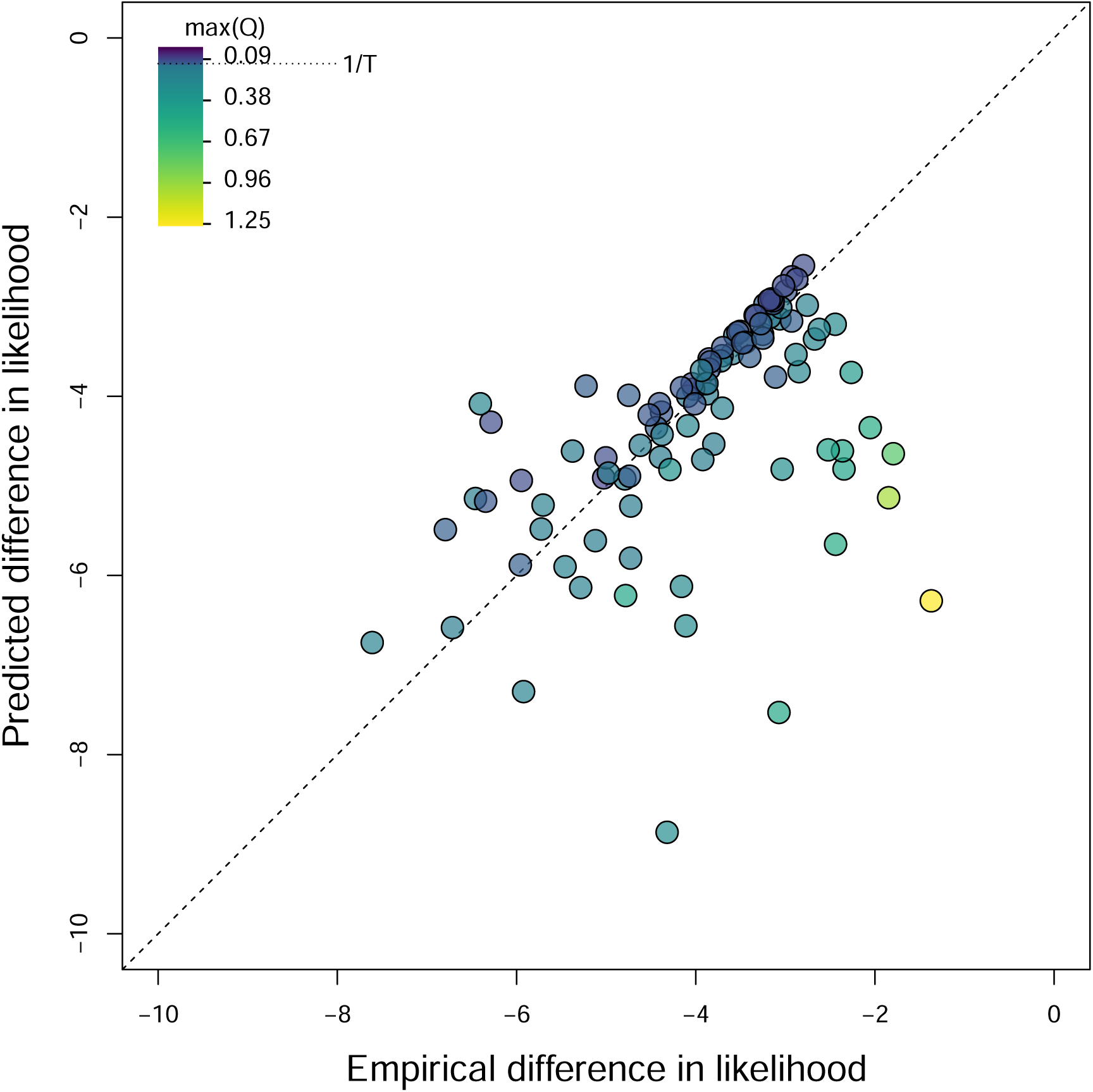
Darwin’s scenario–the singular origin of two coextensive traits on the phylogeny–represents a boundary case to finding the correlation between discrete characters. Pagel’s correlation test for Darwin’s scenario can essentially be reduced to the difference in probability between choosing the same branch twice vs. choosing the branch only once. We demonstrate that here, showing our predicted differences in log likelihood between the independent and dependent trait models (y-axis) against the empirical estimates of the difference in log likelihood between models for simulated Darwin’s scenarios on different phylogenies. Dotted line indicates equality. Points falling off the line represent slight violations of the assumptions we used to derive our prediction. Particularly, we assume that the rates of gain of the traits are so low that only one shift is ever observed. The color of the points indicates cases where this assumption is violated, as outlying points with max(Q) values much greater than 1/*T* (the value of *q*_01_ at which exactly 1 shift is expected) are much more likely to fall off the predicted line. This figure differs from Figure 6 in the main text in that estimated likelihoods are not constrained to fit the assumptions we used to derive the predicted difference in likelihood. Specifically, we do not assume irreversibility, we allow partial correlations, and we do not constrain gain rates to be equal for the independent case.

A similar procedure can be performed if the network is instead what is found in Figure 9B. Here, we simulate the liability *M*_*liab*_ by Brownian Motion (root = 0, *σ*^2^ = 1). Body size (B) is then a function of this liability using the reciprocal equation, *B* = *M*_*liab*_ + *ϵ*. We then discretize migratory behavior (M) from the liability as before, and simulate values of abundance using the same equation *N* = *B* + *-*3*M* + *ϵ*.

We acknowledge that the manner in which the data is generated is somewhat contrived and parameters were chosen to produce a figure that maps on to the familiar conceptual depiction of Simpson’s paradox. This was done to aid visual and conceptual interpretation. Under a wider range of parameter combinations, such consistent differences between PGLS and OLS results will often break down — particularly due to the lack of robustness of PGLS results to violations of Brownian Motion and its sensitivity to singular events (which will often result from our imposition of a threshold model, see Case Study I in the main text). Thus, our primary goal in generating this figure was to choose parameter sets and networks that visually illustrated the phenomenon of Simpson’s paradox in phylogenetic comparative datasets in a clearly interpretable way without substantially violating the assumptions of PGLS regression. While other parameter sets will produce less visually obvious results, the key points of our argument will remain unchanged.

